# A Three Dimensional Immunolabeling Method with Peroxidase-fused Nanobodies and Fluorochromized Tyramide-Glucose Oxidase Signal Amplification

**DOI:** 10.1101/2024.10.25.620157

**Authors:** Kenta Yamauchi, Masato Koike, Hiroyuki Hioki

## Abstract

Three dimensional immunohistochemistry (3D-IHC), immunolabeling of 3D tissues, reveals the spatial organization of molecular and cellular assemblies in the context of the tissue architecture. Deep and rapid penetration of antibodies into 3D tissues and highly sensitive detection are critical for high-throughput imaging analysis of immunolabeled 3D tissues. Here, we report a nanobody (nAb)-based 3D-IHC, POD-nAb/FT-GO 3D-IHC, for high-speed and high-sensitive detection of targets within 3D tissues. Peroxidase-fused nAbs (POD-nAbs) enhanced immunolabeling depth and allowed for highly sensitive detection by combined with a fluorescent tyramide signal amplification system, Fluorochromized Tyramide-Glucose Oxidase (FT-GO). Multiplex labeling was implemented to the 3D-IHC by quenching POD with sodium azide. Using the 3D-IHC technique, we successfully visualized somata and processes of neuronal and glial cells in millimeter-thick mouse brain tissues within three days. Given its high-speed and high-sensitive detection, our 3D-IHC protocol, POD-nAb/FT-GO 3D-IHC, would provide a useful platform for histochemical analysis in 3D tissues.

## Introduction

Three dimensional immunohistochemistry (3D-IHC), immunolabeling of 3D tissues, visualizes spatial cellular organization and molecular distributions within large-scale biological and clinical tissues. 3D-IHC provides rich cellular and molecular information and permits reconstruction of microstructural details, unbiased sampling throughout a tissue, and a high data throughput, revealing molecule-structure-function relationships in a biological system. Recent advances in 3D-IHC methods combined with tissue clearing techniques and advanced light microscopy have allowed for visualizing molecular expression within organoids, organs, murine and human embryos, whole mice and even human organs^1–6^. Implementation of 3D-IHC to clinical tissues would facilitate more accurate and objective pathological diagnosis^7,8^.

Deep penetration of antibodies (Abs) into 3D tissues is required for high staining homogeneity in 3D-IHC. The insufficient depth of Abs in 3D-IHC typically results in the peripheral deposition of Abs in 3D tissues, leading to a large gradient of immunosignals. Several approaches have been designed to improve penetration depth of Abs and staining homogeneity^9^. For example, Ab diffusion is facilitated by tissue permeabilization^1,5,10–12^ and Ab incubation at high temperature^13^. Transient inhibition of antigen–antibody binding by manipulation of temperature, pH, and ionic strength is adapted to enhance Ab penetration^1,13–16^. High pressure transcardial perfusion of Abs and tissue compression following to transformation into elastic hydrogels reduce physical distances for Abs to diffuse to reach deep region during immunolabeling^6,17,18^. Additionally, active Ab transport, such as electrophoresis and pressure, have been used on acrylamide-embedded tissues^15,19^. However, despite these technical advances, insufficient penetration of conventional Abs, such as immunoglobulin G (IgG), M (IgM), and Y (IgY), has remained an insuperable barrier for immunolabeling of 3D tissues.

Nanobodies (nAbs) or variable domain of heavy chain of heavy chain antibodies (VHH antibodies), recombinant minimal antigen binding fragments from single-chain Abs in camelids and selachians, have been used for research, diagnostic and therapeutic reagents^20^. NAbs should be particularly suitable for 3D immunolabeling due to their much smaller in size (12–15 kDa) than conventional IgG Abs (∼150 kDa). Indeed, nAbs have been shown deep penetration into biological tissues compared to conventional antibodies^3,21–23^ and further implemented to whole-body immunolabeling by homogenous delivery of them with high-pressure active perfusion^3,24^. The major issue in the implementation of nAbs to 3D-IHC is signal strength. While the signal in immunofluorescence (IF) with conventional Abs including IgG, IgM and IgY, is typically amplified by secondary IgG Abs that tolerate many labels per molecule and bind to distinct epitopes of a primary Ab, nAbs are conjugated with one or two synthetic fluorophores and used for IF without signal amplification. Signal strength imposes a limit on the throughput of imaging of 3D-IHC, where much long image acquisition time is required.

Tyramide signal amplification (TSA) might provide a solution to the relatively low sensitivity of 3D-IHC using nAbs. TSA is a highly sensitive enzymatic amplification method that allows detection of low-abundance target and dramatic signal enhancement. TSA involves the catalytic activity of peroxidase (POD) to yield high-density labeling of targets^25,26^. TSA was originally developed for the detection of immunosorbent and immunoblotting assays^25^. Soon after its development, TSA was widely adapted to histochemical analysis, which includes IHC^27^, DNA and RNA *in situ* hybridization (ISH)^28–30^, and electron microscopy^31,32^. In our earlier studies, we reported straightforward and cost-effective TSA systems, namely, Biotinyl Tyramine-Glucose Oxidase (BT-GO)^33,34^ and Fluorochromized Tyramide-Glucose Oxidase (FT-GO)^35^. Unlike conventional ones, these TSA systems utilize hydrogen peroxide (H_2_O_2_) produced by oxidation of glucose by glucose oxidase^33–35^. Enzymatic reaction between glucose and glucose oxidase supplies H_2_O_2_ stably during their reaction to improve operational stability of these TSA systems.

Here, we developed an nAb-based 3D immunolabeling, namely POD-nAb/FT-GO 3D-IHC, for high-speed and high-sensitive detection of target molecules within 3D tissues. To this end, we combined POD-fused camelid nAbs (POD-nAbs or P-RAN-bodies^36^) with our original fluorescent TSA system, FT-GO^35^. POD-nAb/FT-GO 3D-IHC allows for visualization of target molecules in mouse brain slices of 1-mm thickness with drastic signal enhancement within three days. For multiplex labeling in the immunolabeling of 3D tissues, POD-nAbs were applied sequentially after quenching nAb-fused POD with high concentration of sodium azide (NaN_3_). Using the 3D-IHC technique, we visualized somata and processes of neurons and glia in millimeter-thick mouse brain tissues within three days. We further showed microglial activation in thick brain slices of an Alzheimer’s disease (AD) model mouse with respect to ß-amyloid plaques (Aß plaques).

## Results

### POD-nAb/FT-GO 3D-IHC

For high-speed and high-sensitive detection of targets within large-scale tissues, we built an nAb-based volume immunolabeling, POD-nAb/FT-GO 3D-IHC (Fig. 1). POD-nAb/FT-GO 3D-IHC incorporates three main components: (1) tissue permeabilization with Sca*l*eA2 solution^37^, (2) POD-nAb binding to their targets and (3) FT-GO reaction within large-scale tissues. POD-nAb (P-RAN-body) is a genetically encoded immunoreagent, which consists of a camelid nAb and a variant of horseradish peroxidase (HRP)^36^ (Fig. 1a). The culture medium of mammalian cell line 293T cells transfected with a POD-nAb expression vector is used for immunolabeling. The staining procedure of POD-nAb/FT-GO 3D-IHC is based on AbSca*l*e method^11^. For sensitive detection of POD-nAbs binding to their respective antigens, a multiplex fluorescent TSA signal amplification system, FT-GO, is implemented to the 3D-IHC method. POD-nAb/FT-GO 3D-IHC can visualize the spatial organization of target molecules in millimeter-thick mouse brain tissues within three days. Following an incubation in Sca*l*eA2 solution for 24 hr, mouse brain tissues are reacted with POD-nAbs for 20-24 hr. Using a TSA reaction with peroxidase activity of POD-nAbs, FT is deposited onto brain tissues within 8.5 hr (Fig. 1b). Sca*l*eA2 treatment can be omitted for preservation of antigenicity. Figure 1c shows an example of POD-nAb/FT-GO 3D-IHC. A 1-mm-thick mouse brain slice infected with an AAV2/PHP.eB CAG-EGFP-WPRE vector is subjected to POD-nAb/FT-GO 3D-IHC. A POD-nAb against GFP is used for the 3D immunolabeling. Maximum intensity projection (MIP) images of the 1-mm-thick mouse brain slice are represented (Fig. 1c). Somata and processes of neurons and astrocytes in a millimeter-thick brain slice are clearly and intensely labeled by POD-nAb/FT-GO 3D-IHC method within three days.

**Figure 1.**
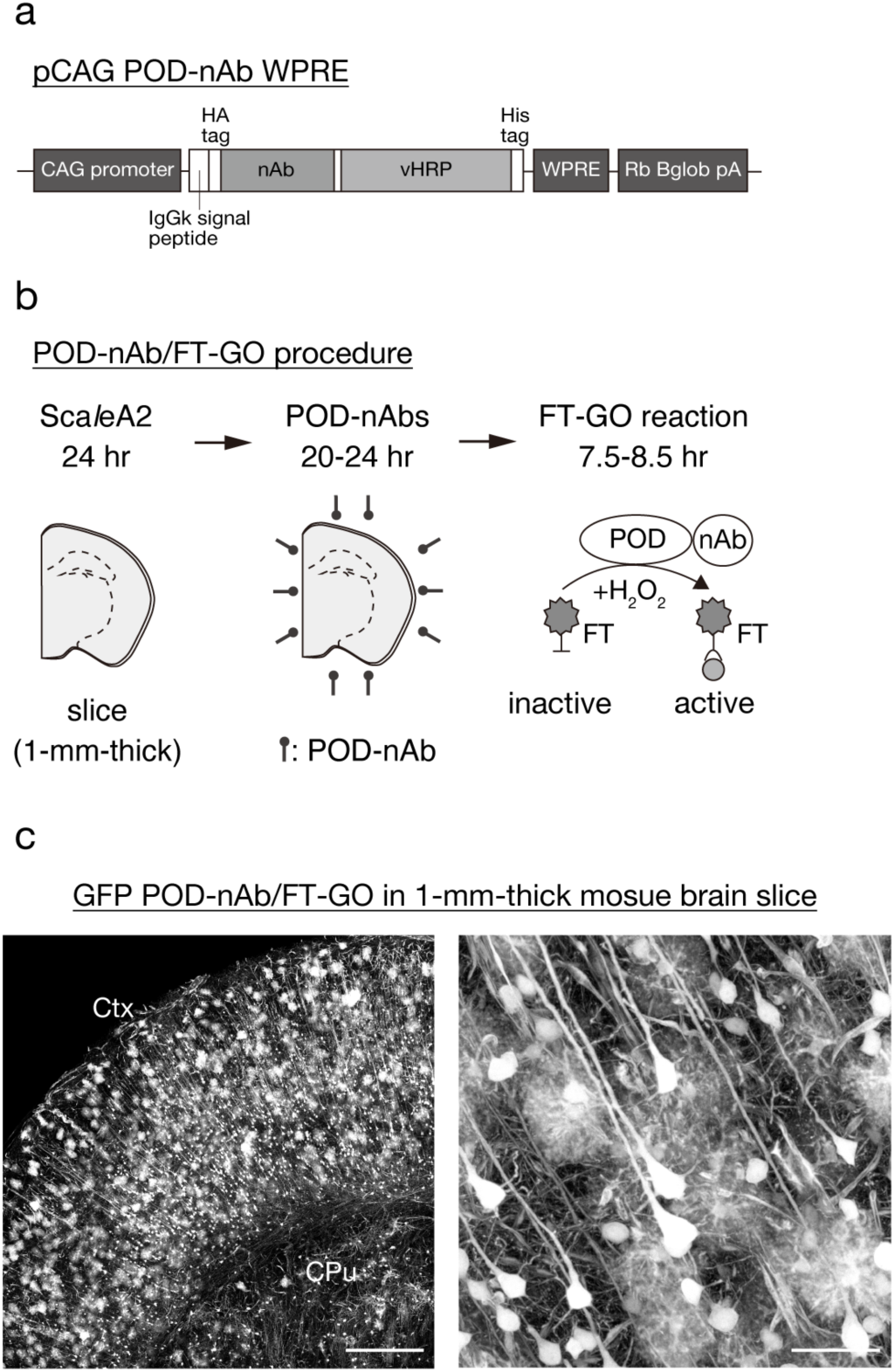
POD-nAb/FT-GO 3D-IHC. **a**) The vector structure of pCAG POD-nAb WPRE. POD-nAbs are camelid nAbs fused with POD. **b**) The schedule and procedure for POD-nAb/FT-GO 3D-IHC. **c**) GFP POD-nAb/FT-CO 3D-IHC in a 1-mm-thick mouse brain slice infected with AAV2/PHP.eB CAG-EGFP-WPRE. Maximum intensity projection (MIP) images of the 1-mm-thick brain slice are represented. The right panel shows a higher magnification image of the left panel. CPu: caudate-putamen, Ctx: cerabral cortex, FT: fluorochromized tyramide, Rb Bglob pA: rabbit beta-globin polyadenylation signal, vHRP: an enhanced variant of horseradish peroxidase, WPRE: woodchuck hepatitis virus posttranscriptional regulatory element. Scale bar: 500 µm in (**c, left**) and 50 µm in (**c, right**).

### Superior tissue penetration of POD-nAbs

We first tested tissue permeability of POD-nAbs and conventional IgG and IgY Abs into mouse brain slices of 1-mm thickness (Fig. 2). POD-nAbs are nAbs fused with a variant of HRP and their molecular weights are roughly 60 kDa, which is approximately 60% smaller in size compared with conventional IgG Abs (∼150 kDa). Given that the diffusivity of a molecule in a fluid medium is inversely related to its molecular weight, the smaller sizes of POD-nAbs should enhance tissue penetration. To assess tissue permeability of immunoreagents, conventional Abs and POD-nAbs against GFP or RFP were reacted with 1-mm-thick mouse brain slices expressing EGFP or tdTomato for four days. Tissue sections of 40-µm thickness were cut perpendicularly to the brain slices (re-sectioning) and reacted with a fluorophore-conjugated anti-HRP Ab or secondary Abs against IgG or IgY of the host species (Fig. 2a). Two polyclonal Abs and one monoclonal Ab from different host species were used for each target. AAV-PHP.eB vectors carrying EGFP or tdTomato, a variant of RFP, were intravenously administrated for brain-wide expression of the fluorescent proteins (FPs)^38^. Strikingly, POD-nAbs penetrated deep inside mouse brain tissues (Fig. 2b-i). While immunoreactivities of conventional Abs against GFP were only restricted on the surface of brain slices (Fig. 2c-e), a GFP POD-nAb penetrated deep and labeled EGFP-positive cells located at the center of 1-mm-thick brain slices (Fig. 2b). The uniform signal distribution irrespective of depth from the surface was also found in immunolabeling with an RFP POD-nAb (Fig. 2f). All of tested conventional IgG Abs against RFP failed to penetrated deep enough to reach the center of slices (Fig. 2g-i).

**Figure 2.**
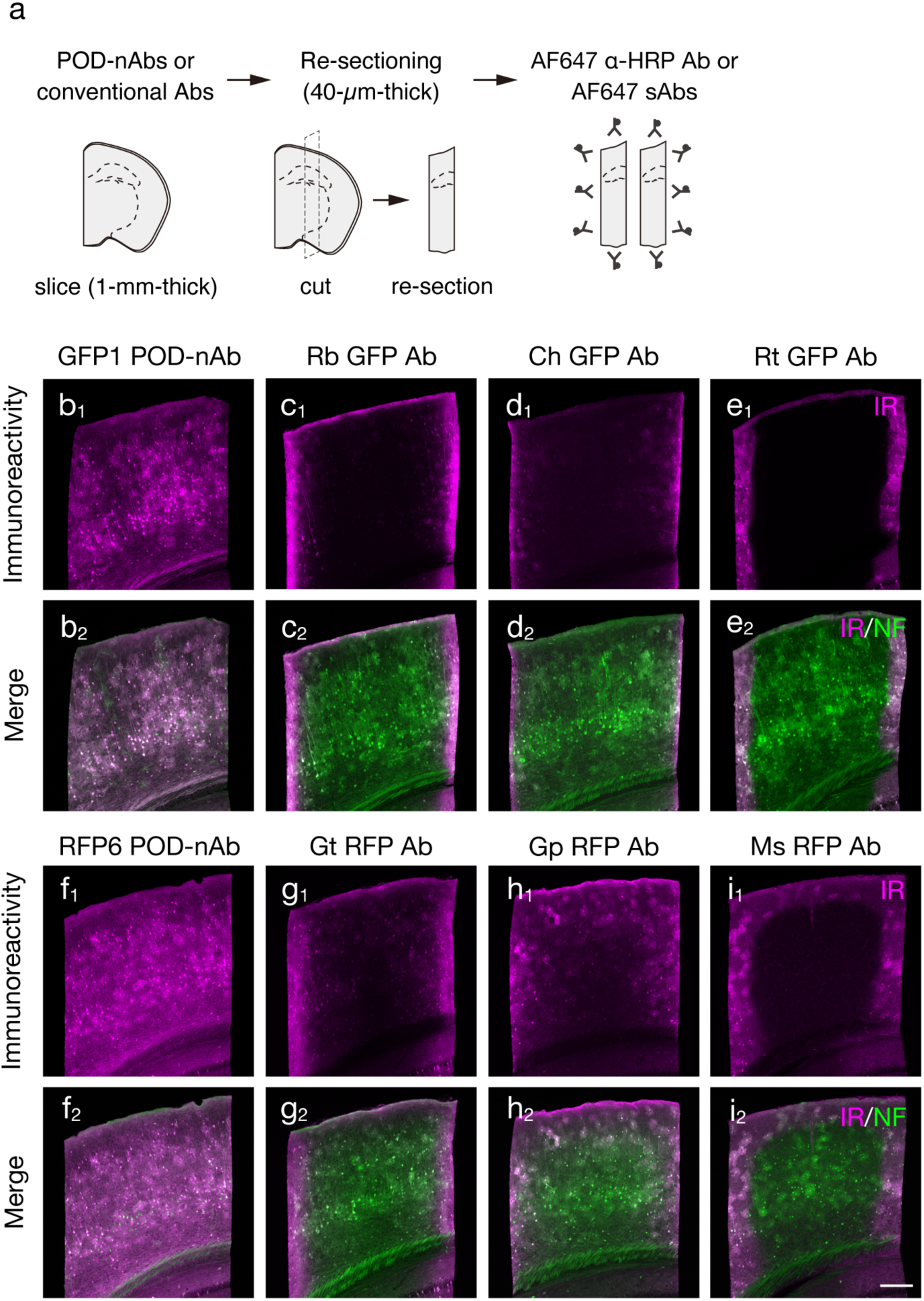
Superior tissue penetration of POD-nAb. **a**) Schematic diagram of an experimental procedure for testing tissue penetration of POD-nAbs and conventional Abs. **b-e**) Tissue penetration of a POD-nAb (**b**) and conventional Abs against GFP (**c-e**) (n = 3 animals for each condition). 1-mm-thick brain slices infected with AAV2/PHP.eB CAG-EGFP-WPRE are reacted with a POD-nAb (**b**), rabbit polyclonal Ab (**c**), chicken polyclonal Ab (**d**), and rat monoclonal Ab (**e**) against GFP. **b_1_, c_1_, d_1_, e_1_**) Representative images of immunoreactivity (IR) for GFP Abs in the cerebral cortex. **b_2_, c_2_, d_2_, e_2_**) Merged images of IR for GFP (magenta) and EGFP fluorescence (green) in the cerebral cortex. **f-i**) Tissue penetration of a POD-nAb (**f**) and conventional Abs against RFP (**g-i**) (n = 3 animals for each condition). 1-mm-thick brain slices infected with AAV2/PHP.eB CAG-tdTomato-WPRE are reacted with a POD-nAb (**f**), goat polyclonal Ab (**g**), giunea pig polyclonal Ab (**h**), and mouse monoclonal Ab (**i**) against RFP. **f_1_, g_1_, h_1_, i_1_**) Representative images of IR for RFP Abs in the cerebral cortex. **f_2_, g_2_, h_2_, i_2_**) Merged images of IR for RFP (magenta) and tdTomato fluorescence (green) in the cerebral cortex. AF: Alexa Fluor, IR: immunoreactivity, NF: native fluorescence. Scale bar: 200 µm.

Tissue permeabilization protocols have been developed for volume labeling to enhance tissue penetration of chemical dyes and Abs. These include denaturing agent-based^11^, detergent-based^2,12^, hydrogel-based^10,39^ and organic solvent-based^1,40^ protocols. Additionally, tissue permeabilization combining components of different protocols have been implemented to several 3D-IHC techniques^5,41–43^. Tissue permeabilization protocols for volume labeling can affect immunoreactivity of Abs^1^. We thus examined effects of tissue permeabilization protocols on immunoreactivity of GFP and RFP POD-nAbs (Supplementary Figure 1, 2). We choose Sca*l*e, CUBIC-HistoVIsion, PACT and iDISCO as denaturing agents-, detergent-, hydrogel-and organic solvent-based protocols, respectively. Following to tissue permeabilization, 1-mm thick brain slices expressing EGFP or tdTomato were cut into 40-µm-thick brain sections and reacted with GFP or RFP POD-Abs. The reacted POD-nAbs were visualized with a fluorophore-conjugated Ab against HRP (Supplementary Figure 1a, 2a). Immunoreactivity of the GFP and RFP POD-nAbs was preserved in re-sections treated with AbSca*l*e and CUBIC-HistoVision protocols (Supplementary Figure 1b-d, 2b-d). In contrast, the immunoreactivity was markedly reduced following to tissue permeabilization by PACT and iDISCO protocols (Supplementary Figure 1b_1_, e_1_, f_1_, 2b_1_, e_1_, f_1_). Additionally, EGFP and tdTomato fluorescence were dramatically quenched by these tissue permeabilization protocols (Supplementary Figure 1b_2_, e_2_, f_2_, 2b_2_, e_2_, f_2_). Given Sca*l*eS tissue clearing techniques show superior tissue integrity applicable to electron microscopic analysis^11,44^, we implemented Sca*l*e protocols for tissue permeabilization in our volume immunolabeling with POD-nAbs. We then asked whether Sca*l*e tissue permeabilization protocol, an incubation in Sca*l*eA2 solution for 24 hr at 37 °C, enhanced immunolabeling depth of POD-nAbs (Fig. 3). 1-mm-thick brain slices infected with AAV-PHP.eB CAG-EGFP or tdTomato-WPRE were reacted with GFP or RFP POD-nAbs for 24 hr, respectively with or without Sca*l*eA2 treatment. Tissue penetration of POD-nAbs was markedly enhanced by Sca*l*e tissue permeabilization: while GFP and RFP POD-nAbs failed to reach the center of 1-mm-thick brain slices without Sca*l*e tissue permeabilization (Fig. 3b, d), these POD-nAbs penetrated deep inside brain tissues and labeled FP-positive cells at the center of slices following to Sca*l*e tissue permeabilization (Fig. 3c, e).

**Figure 3.**
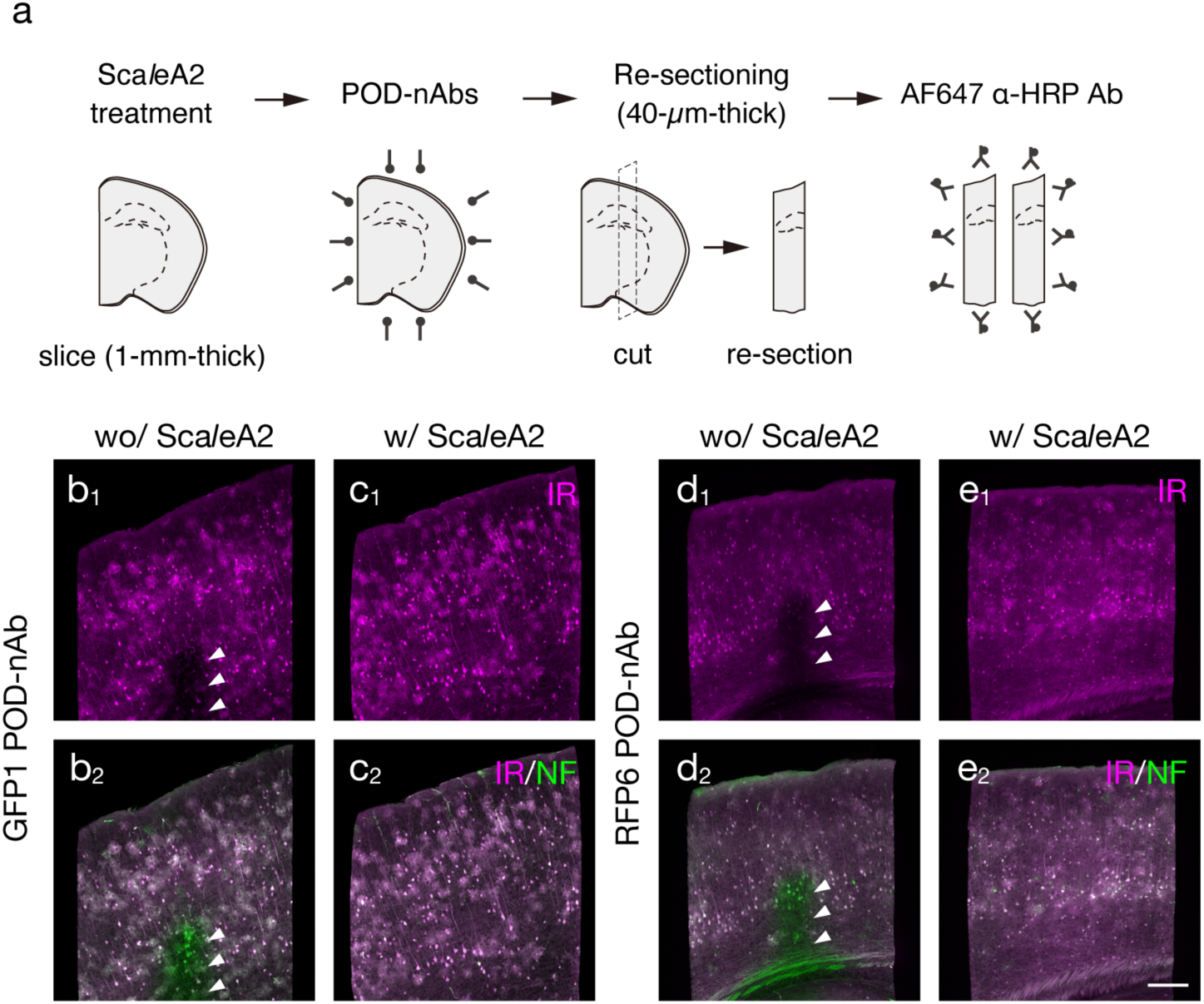
Enhanced penetration of POD-nAbs by Sca*l*eA2 treatment. **a**) Schematic diagram of an experimental procedure for testing Sca*l*eA2 treatment on tissue penetration of POD-nAbs. **b-e)** Tissue penetration of GFP1 (**b, c**) and RFP6 POD-nAb (**d, e**) with (**c, e**) or without (**b, d**) Sca*l*eA2 treatment (n = 3 animals for each condition). Brain slices are infected with AAV2/PHP.eB CAG-EGFP-WPRE (**b, c**) or CAG-tdTomato-WPRE (**d, e**). **b_1_, c_1_, d_1_, e_1_**) Representative images of immunoreactivity (IR) for GFP1 (**b_1_, c_1_**) and RFP6 POD-nAbs (**d_1_, e_1_**) in the cerebral cortex. **b_2_, c_2_, d_2_, e_2_**) Merged images of IR for POD-nAbs (magenta) and EGFP (**b_2_, c_2_**) or tdTomato (**d_2_, e_2_**) fluorescence (green) in the cerebral cortex. Images are acquired with the same parameters for comparisons. Arrowheads indicate insufficient penetration of POD-nAbs. AF: Alexa Fluor, IR: immunoreactivity, NF: native fluorescence. Scale bar: 200 µm.

### Signal amplification with POD-nAb/FT-GO in 3D imaging

TSA is a highly sensitive enzymatic method that enables dramatic signal enhancement in histochemical analysis. TSA utilizes POD to catalyze covalent deposition of haptenized and fluorochromized tyramides in the vicinity of the immobilized enzyme^25,26^. Recently, we developed FT-GO as a multiplex fluorescent TSA system^35^. FT-GO involves POD-catalyzed deposition of FT using H_2_O_2_ produced by oxidation of glucose by glucose oxidase and yields approximately 10-to 30-fold signal amplification compared with indirect IF detections^35^. Strong fluorescence signal obtained using FT-GO signal amplification could deliver a significant enhancement of throughput of imaging of large-scale tissues. We thus examined applicability and scalability of FT-GO signal amplification to 3D-IHC with POD-nAbs (Fig. 4). Following to Sca*l*e tissue permeabilization and reaction with POD-nAbs against GFP or RFP, 1-mm-thick brain slices infected with AAV-PHP.eB CAG-EGFP or tdTomato-WPRE were incubated in an FT-GO reaction mixture containing CF647 tyramide. FT-GO reaction was then initiated by addition of β-D-glucose into the reaction mixture. After the reaction and re-sectioning, tissue sections were subjected to confocal laser scanning microscopy (CLSM) (Fig. 4a). Fluorescence signals obtained by FT-GO wihtin 1-mm-thick brain slices were well matched with EGFP and tdTomato fluorescence in the re-sections (Fig. 4b, c). Quantitative analysis demonstrated FT depositions in 98.8 ± 1.0% EGFP-and 98.7 ± 0.6% tdTomato-positive neurons (Fig. 4d, e). FT-GO signal amplification following to POD-nAbs incubation (POD-nAb/FT-GO reaction) showed superior sensitivity compared with of FPs: the number of FT-labeled neurons was larger than that of FP-labeled neurons in the re-sections (GFP-positive neurons in FT-positive neurons: 90.0 ± 2.3%, tdTomato-positive neurons in FT-positive neurons: 85.7 ± 4.3%; Fig. 4d, e).

**Figure 4.**
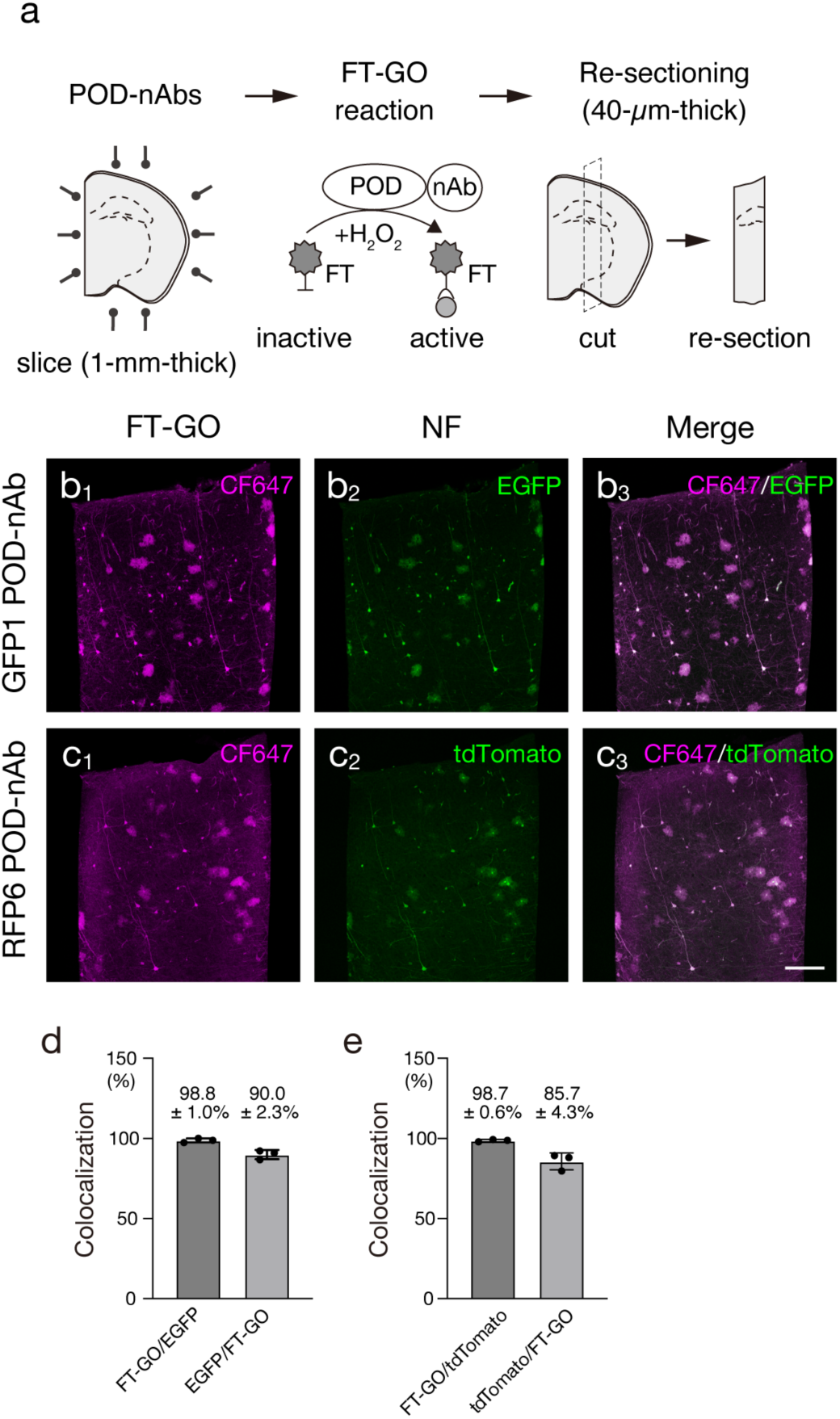
3D-TSA reaction with FT-GO. **a**) Schematic diagram of an experimental procedure for testing 3D-TSA reaction with FT-GO and POD-nAbs. **b, c**) 3D FT-GO reaction in 1-mm-thick brain slices with GFP1 POD-nAb (**b**) and RFP6 POD-nAb (**c**) (n = 3 animals for each condition). 1-mm-thick brain slices are infected with AAV2/PHP.eB CAG-EGFP-WPRE (**b**) or CAG-tdTomato-WPRE (**c**). **b_1_, _2_, c_1_, _2_**) Representative images of CF647-tyramide deposition with GFP1 POD-nAb (magenta, **b_1_**) and RFP6 POD-nAb (magenta, **c_1_**), and EGFP (green, **b_2_**) and tdTomato (green, **c_2_**) fluorescence in the cerebral cortex. **b_3_, c_3_**) Merged images of the CF647-tyramide deposition (magenta) and EGFP (**b_3_**) or tdTomato (**c_3_**) fluorescence (green) in the cerebral cortex. **d, e**) Histograms showing the percentages of FT-GO^+^ cells in EGFP^+^ cells and EGFP^+^ cells in FT-GO^+^ cells (n = 466 cells, EGFP^+^ cells; n = 507 cells, FT-GO^+^ cells from 3 animals) (**d**), and FT-GO^+^ cells in tdTomato^+^ cells and tdTomato^+^ cells in FT-GO^+^ cells (n = 619 cells, tdTomato^+^ cells; n = 704 cells, FT-GO^+^ cells from 3 animals) (**e**). Data are represented as means ± SDs. FT: fluorochromized tyramide, NF: native fluorescence. Scale bar: 200 µm.

FT-GO allows signal enhancement in IF on tissue sections^35^. We thus explored signal amplification characteristics of FT-GO in 3D immunolabeling with POD-nAbs. 1-mm-thick mouse brain slices infected with an AAV2/PHP.eB CAG-EGFP-WPRE vector were subjected to POD-nAb/FT-GO reaction. Following to the reaction, the brain slices were cleared with Sca*l*eS4 solution^11^, mounted in a 3D-printed chamber, and imaged by CLSM^44–46^ (Fig. 5a). For visualization of EGFP fluorescence, 1-mm-thick brain slices infected with the AAV-PHP.eB vector carrying EGFP were cleared with Sca*l*eSF method^44^ (Fig. 5a). Critically, fluorescence signals obtained by POD-nAb/FT-GO reaction was much stronger than those of EGFP (Fig. 5b): while POD-nAb/FT-GO reaction clearly visualized somata and processes of neuronal and glial cells, only faintly labeled cells were observed with EGFP fluorescence under the same imaging condition. POD-nAb/FT-GO reaction yielded 9.0-fold signal amplification at the depth of 500 µm below the surface of slices compared with EGFP fluorescence (Fig. 5c). Synthetic fluorophore conjugated nAbs penetrate deep inside tissues and provide a contrast to overcome tissue autofluorescence in 3D-IHC^3,24^. Indeed, a synthetic fluorophore conjugated nAb against GFP showed homogenous labeling in 1-mm-thick brain slices infected with the AAV-EGFP vector (Supplementary Figure 3). We then compared fluorescent signals obtained by direct IF with a synthetic fluorophore conjugated nAb with those by POD-nAb/FT-GO reaction. After tissue permeabilization using Sca*l*eS protocol, the synthetic fluorophore conjugated nAb against GFP was reacted with 1-mm-thick brain slices infected with AAV2/PHP.eB CAG-EGFP-WPRE. The brain slices were subjected to tissue clearing with Sca*l*eS4 solution followed by CLSM (Fig. 5a). POD-nAb/FT-GO reaction provided dramatically increased sensitivity in 1-mm-thick brains slices, compared with IF using an nAb conjugated with a synthetic fluorophore (Fig. 5d). Fluorescent signals obtained by POD-nAb/FT-GO reaction were 6.8-fold stronger than those by direct IF with the synthetic fluorophore conjugated nAb at the depth of 500 µm below the surface of slices (Fig. 5e).

**Figure 5.**
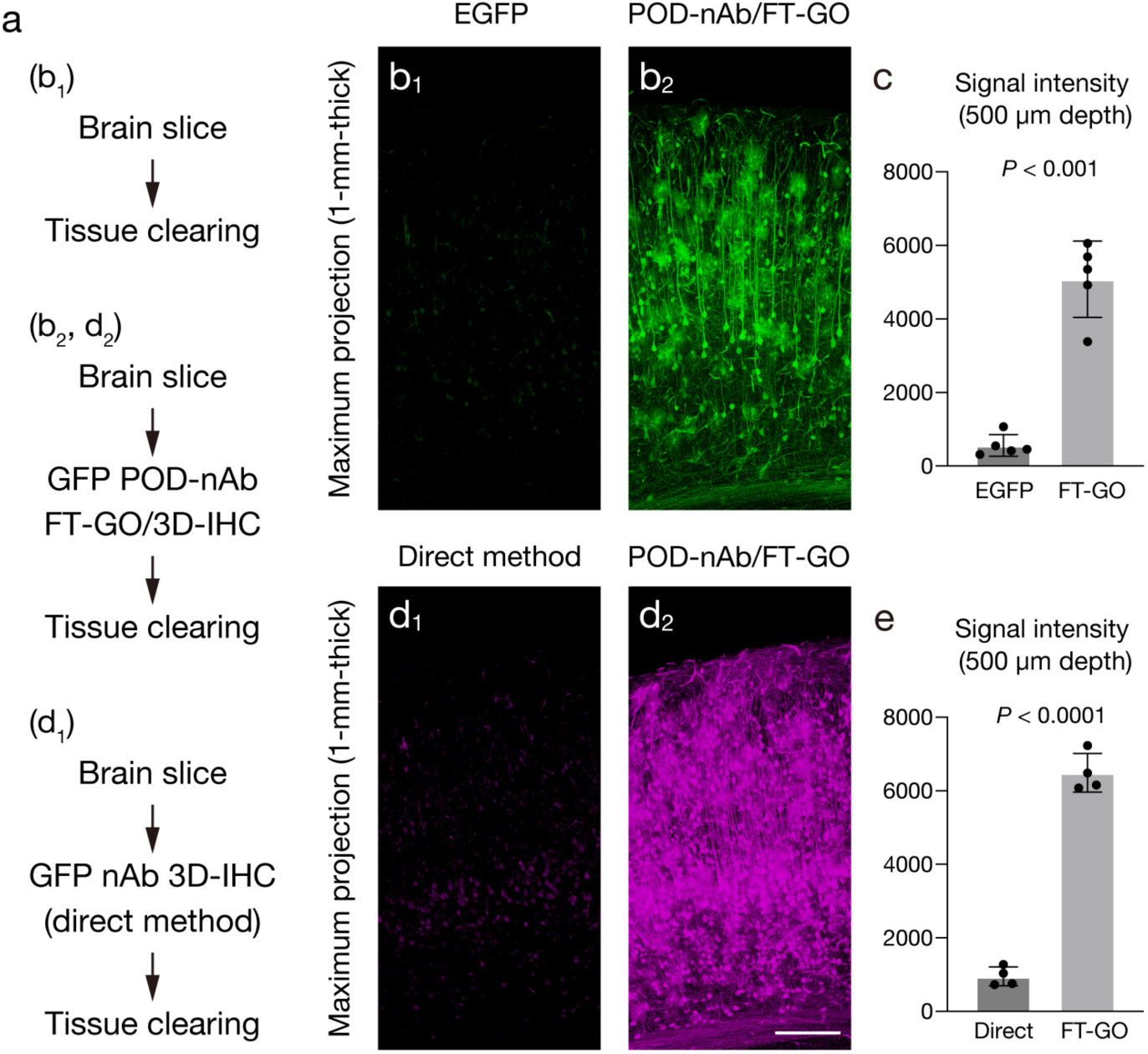
Signal amplification with POD-nAb/FT-GO in 3D imaging. **a**) Experimental procedures in each panel. Following tissue clearing, brain slices of 1-mm thickness are subjected to CLSM imaging. **b**) Maximum intensity projection (MIP) images from EGFP fluorescence (**b**_1_) and POD-nAb/FT-GO 3D-IHC (**b_2_**) in the cerebral cortex of 1-mm-thick brain slices (n = 5 animals for each condition). Brain slices are infected with AAV2/PHP.eB CAG-EGFP-WPRE. CF488A tyramide is used for color development. Images are acquired with the same parameters for comparisons. **c**) Histograms representing relative fluorescence intensity at the depth of 500 µm (n = 588 cells from 5 animals, EGFP fluorescence; n = 520 cells from 5 animals, POD-nAb/FT-GO 3D-IHC; *t* = 9.360, *df* = 4.644, *P* = 0.0003, Welch’s t test). **d**) MIP images from 3D-IHC with an Alexa Fluor 647-conjugated GFP nAb (**d_1_**) and POD-nAb/FT-GO 3D-IHC (**d_2_**) in the cerebral cortex of 1-mm-thick brain slices (n = 4 animals for each condition). Brain slices are infected with AAV2/PHP.eB CAG-EGFP-WPRE. CF647 tyramide is used for color development. Images are acquired with the same parameters for comparisons. **e**) Histograms representing relative fluorescence intensity at the depth of 500 µm (n = 281 cells from 4 animals, GFP nAb 3D-IHC; n = 375 cells from 4 animals, POD-nAb/FT-GO 3D-IHC; *t* = 18.93, *df* = 4.353, *P* < 0.0001, Welch’s t test). NF: native fluorescence. Data are represented as means ± SDs. Scale bar: 200 µm.

### Detection of an endogenous protein with POD-nAb/FT-GO 3D-IHC

We next asked whether POD-nAb/FT-GO 3D-IHC could be implemented to detect an endogenous protein in large-scale tissues. For this, we generated a POD-nAb against integrin alpha M (ITGAM) (also known as cluster of differentiation molecule 11b [CD11b]) using a published sequence^47^. ITGAM is expressed by myeloid lineage cells, including neutrophils, monocytes and macrophages. ITGAM has gained usage as a marker for microglial cells, which serve as resident macrophages within the CNS^48^. Specific labeling of POD-nAb/FT-GO 3D-IHC for ITGAM was tested by FT-GO IF on re-sections with an IgG antibody against allograft inflammatory factor 1 (AIF1) (also known as ionized calcium binding adaptor molecule 1 [Iba1]), a reliable marker for microglia (Fig. 6a). Following to incubation with the ITGAM POD-nAb and FT-GO reaction, mouse brain slices of 1-mm thickness were cut perpendicularly into 40-µm-thick brain sections. After quenching nAb-fused POD with high concentration of NaN_335_, FT-GO IF for AIF1 was carried out on the sections. In the 3D-IHC for ITGAM, 1-mm-thick mouse brain slices were not processed for Sca*l*eS tissue permeabilization prior to the POD-nAb reaction. The uniform signal distribution was found irrespective of depth from the surface in the POD-nAb/FT-GO 3D-IHC (Fig. 6b). The ITGAM POD-nAb penetrated deep inside and deposited FT at the center of the slices. At high magnification, we found tortuous and ramified processes radiating from small cell bodies labeled by POD-nAb/FT-GO 3D-IHC for ITGAM, which is characteristic of resting microglial cells (Fig. 6c_1_). Critically, almost all cells labeled with the ITGAM POD-nAb in 1-mm-thick mouse brain slices were also labeled with an IgG antibody against AIF1 in 40-µm-thick re-sections (Fig. 6c_1-3_).

**Figure 6.**
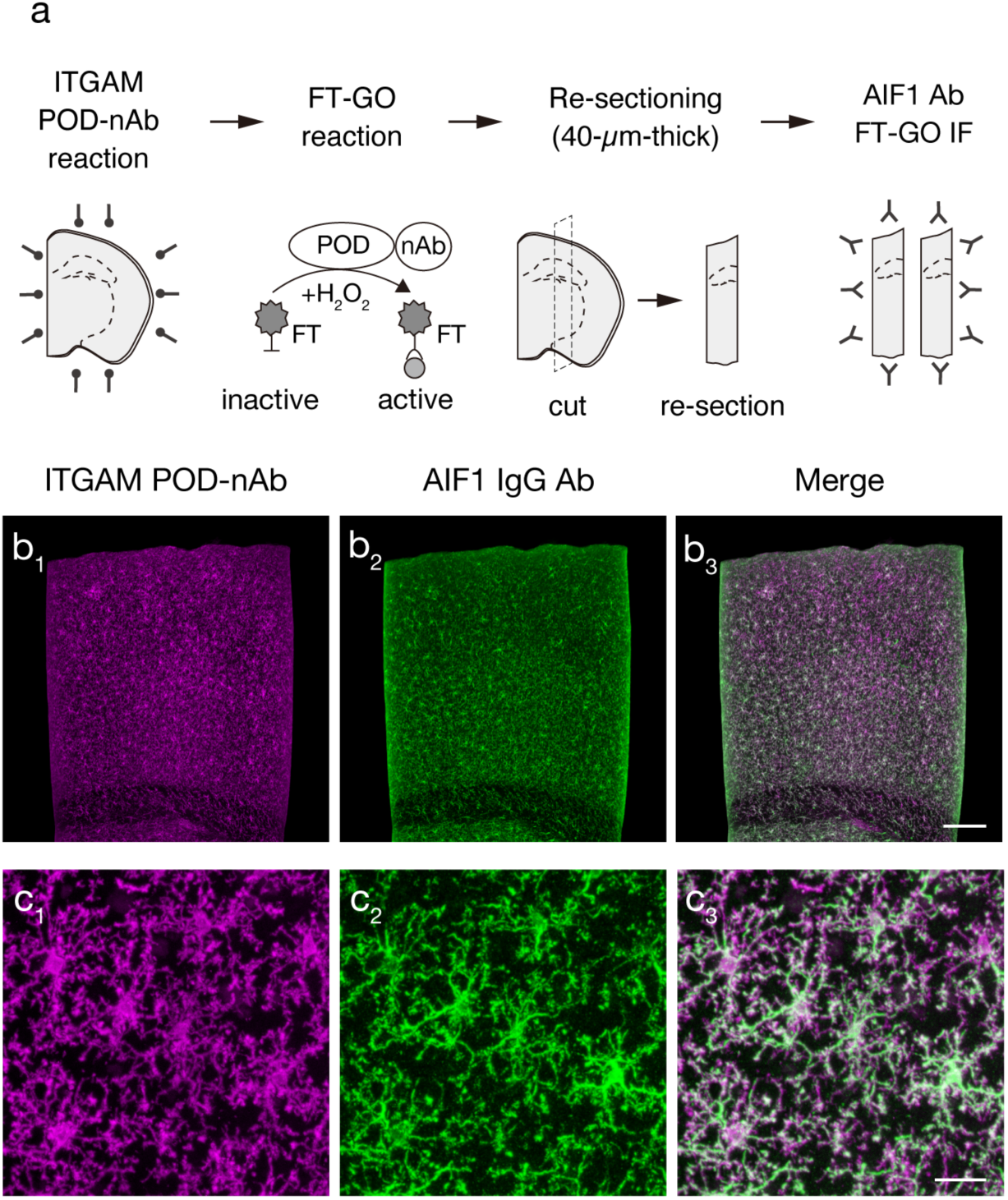
ITGAM POD-nAb/FT-GO 3D-IHC in 1-mm-thick mouse brain slices. **a**) Schematic diagram of an experimental procedure for testing ITGAM POD-nAb/FT-GO 3D-IHC. **b**) AIF1 FT-GO IF in a re-section prepared from a 1-mm-thick brain slices stained by ITGAM POD-nAb/FT-GO 3D-IHC (n = 3 animals). **b_1, 2_**) Representative images of immunoreactivity for a ITGAM POD-nAb (magenta, **b_1_**) and an AIF1 IgG Ab (green, **b_2_**). **b_3_**) A merged image of (**b_1_**) and (**b_2_**). **c**) A high magnification image of (**b**). **c_1, 2_**) Representative images of ITGAM POD-nAb (magenta, **c_1_**) and AIF1 IgG Ab immunoreactivity (green, **c_2_**). **c_3_**) A merged image of (**c_1_**) and (**c_2_**). FT: fluorochromized tyramide. Scale bars: 200 µm in (**b**) and 25 µm in (**c**).

Microglial activation is a histological hallmark in the brain of patients with alzheimer’s disease (AD) as well as mouse models of ß-amyloidosis^49,50^. Microglial cells around Aß plaques undergo morphological change from a ramified to an amoeboid appearance. In parallel with the morphological alternation, they also display different cell surface and intracellular markers from their resting state. Critically, activation of microglial cells leads to an increased expression of ITGAM by them^48^. Microglial activation surrounding Aß plaques is thought to play a detrimental role in the pathogenesis of AD^49,50^. We thus examined applicability POD-nAb/FT-GO 3D-IHC for ITGAM to detection of microglial activation around Aß plaques. We first asked whether the ITGAM POD-nAb can detect local activation of plaque-associated microglia (PAM). For this, FT-GO IF with the ITGAM POD-nAb was carried out on 40-µm-thick brain sections of an *App* knock-in mouse model of ß-amyloidosis, *App^NL-G-F/NL-G-F^* mice^51^ (Supplementary Figure 4). Indeed, increased immunoreactivity for ITGAM, an indicator for microglial activation, was clearly detected in microglia surrounding Aß plaques using the ITGAM POD-nAb (Supplementary Figure 4b_1-4_). We then performed POD-nAb/FT-GO 3D-IHC for ITGAM in 500-µm-thick brain slices of *App^NL-G-F/NL-G-F^* mice. The slices were further treated with 1-Fluoro-2,5-bis(3-carboxy-4-hydroxystyryl)benzene (FSB)^52^ to label amyloid fibrils (Fig. 7a). Critically, POD-nAb/FT-GO 3D-IHC successfully visualized microglial activation with respect to Aß plaques (Fig. 7b). An *xy* image obtained at depths at 250 µm clearly showed activated microglial cells that were intensely labeled with ITGAM POD-nAb in the vicinity of Aß plaques (Fig. 7c). POD-nAb/FT-GO 3D-IHC for ITGAM also labeled resting microglial cells with a ramified morphology apart from Aß plaques (Fig. 7c), demonstrating the high sensitivity of the 3D-IHC protocol.

**Figure 7.**
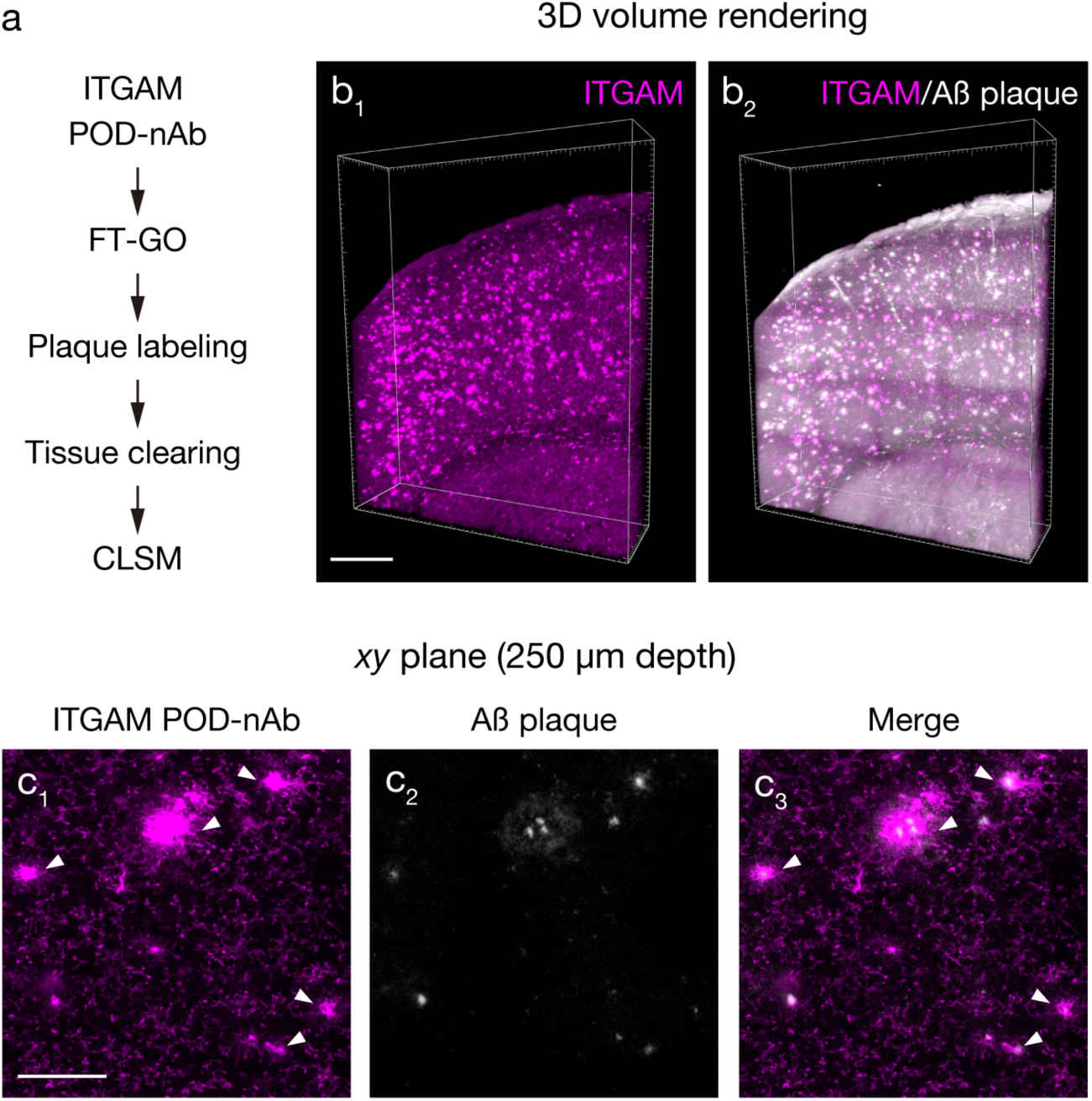
Deep tissue imaging of microglial activation in an AD mouse model. **a**) The procedure of deep tissue imaging of microglial activation with ITGAM POD-nAb/FT-GO 3D-IHC. **b**) 3D volume rendering images of ITGAM POD-nAb/FT-GO 3D-IHC (magenta, **b_1_**) in an *App^NL-G-F/NL-G-F^* mouse brain slice of 500-µm thickness. The slice is also stained with FSB to label amyloid fibrils (n = 3 animals). **b_2_**) A merged volume rendering image of ITGAM (magenta) and FSB (white). **c**) An *xy* images of double labeling with an ITGAM POD-nAb (magenta, **c_1_**) and FSB (white, **c_2_**) labeling in the cerebral cortex of an *App^NL-G-F/NL-G-F^*mouse brain slice at the depth of 250 µm. **c_3_**) A merged images of (**c_1_**) and (**c_2_**). 6 month-old *App^NL-G-F/NL-G-F^*mouse is subjected to the POD-nAb/FT-GO 3D-IHC. Arrowheads indicate activated microglial cells. Scale bar: 500 µm in (**a**) and 100 µm in (**b**).

### Multiplex volume immunolabeling by POD-nAb/FT-GO 3D-IHC

Multiplex volume immunolabeling allows for simultaneous detection of multiple distinct targets within in a single thick tissues or intact organs to analyze the spatial organization of the targets in the context of the tissue architecture^17^. Finally, we demonstrated multiplexing capability of POD-nAb/FT-GO 3D-IHC (Fig. 8). For multiplex labeling with POD-nAb/FT-GO reaction, nAb-fused POD must be inactivated prior to subsequent rounds of detection. NaN_3_ was used for quenching nAb-fused POD activity because of its less deleterious effects on antigenicity, antigen–antibody binding and tissue integrity^35^. Brain slices of 1-mm-thickness were cut from Parvalbumin (PV)/myristoylation-EGFP-low-density lipoprotein receptor C-terminal BAC transgenic mice (PV-FGL mice)^53^ were doubly immunolabeled with the ITGAM and GFP POD-nAbs. In PV-FGL mice, EGFP is specifically expressed and targeted to the somatodendritic plasma membrane of PV-positive neurons. Following to the first round of FT-GO reaction with the ITGAM POD-nAb, the brain slices were treated with high concentration of NaN_3_ [2% (w/v)] to inactivate the nAb-fused POD. The brain slices were then processed for the second round FT-GO reaction with the GFP POD-nAb and cleared with Sca*l*eS4 solution prior to CLSM (Fig. 8a). ITGAM and GFP POD-nAb/FT-GO reaction simultaneously labeled microglial cells and PV^+^ cortical interneurons in 1-mm-thick brain slices of PV-FGL mice (Fig. 8b, c). Specific distribution and no obvious colocalization of ITGAM and GFP immunoreactivities in a projection image from a depth from 437.4 to 469.8 µm demonstrated staining homogeneity and specificity of multiplex labeling with POD-nAb/FT-GO 3D-IHC (Fig. 8d). We further observed highly ramified and thin processes of microglial cells extending to the somatodendritic portion of PV-positive cortical interneurons (Fig. 8d).

**Figure 8.**
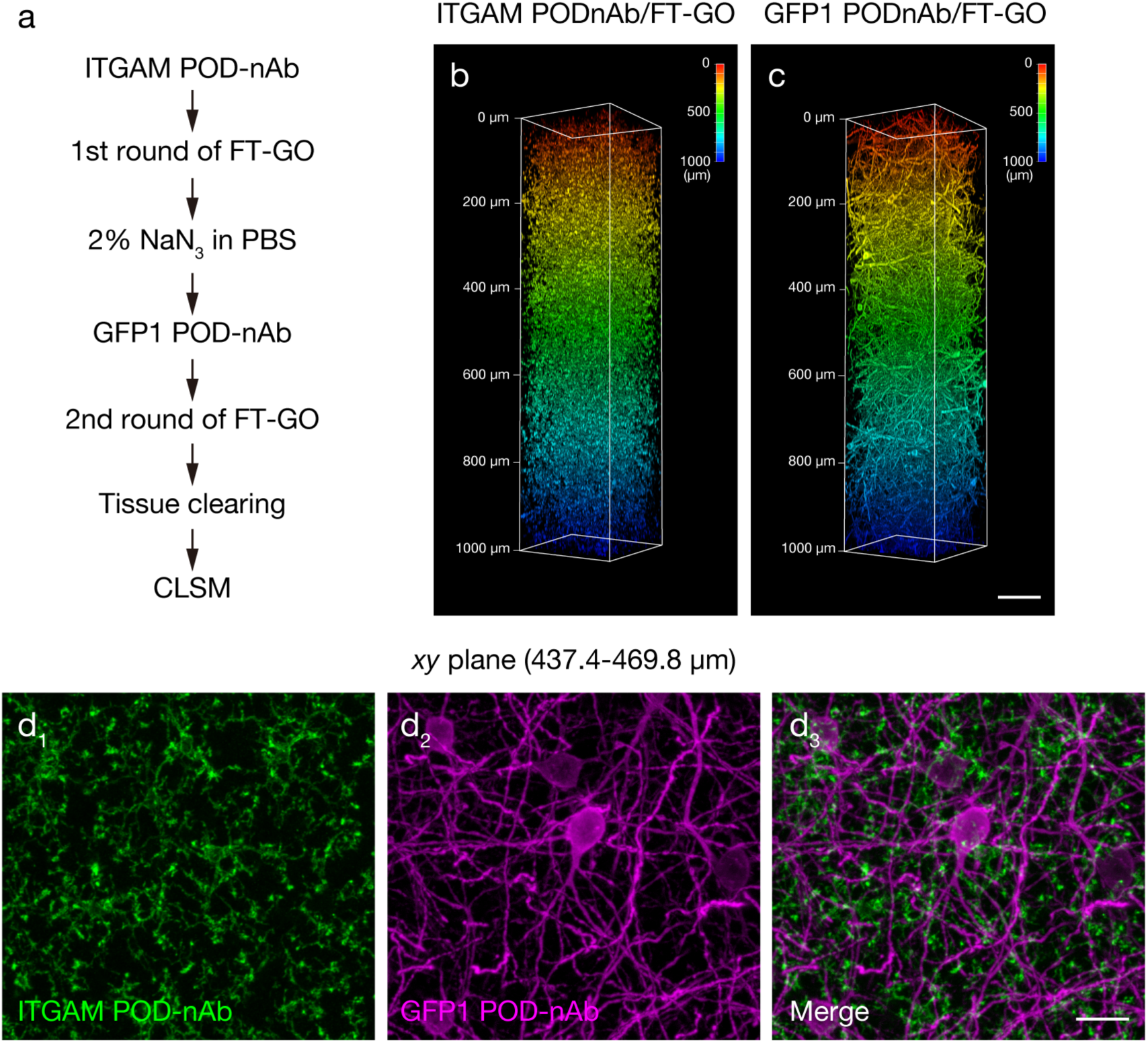
ITGAM and GFP double POD-nAb/FT-GO 3D-IHC in 1-mm-thick brain slices of PV-FGL mice. **a)** The procedure of ITGAM and GFP double POD-nAb/FT-GO 3D-IHC. **b, c**) Depth color coding of 3D volume rendering images of a 1-mm-thick brain slice of a PV-FGL mouse doubly labeled with ITGAM (**b**) and GFP (**c**) POD-nAbs. POD-nAbs are color-developed by 3D FT-GO reaction using CF568 (ITGAM) and CF640R (GFP) tyramides (n = 3 animals). **d**) MIP images of the ITGAM (green, **d_1_**) and GFP (magenta, **d_2_**) double POD-nAb/FT-GO 3D-IHC from a depth of 437.4 to 469.8 µm. **d_3_**) A merged image of (**d_1_**) and (**d_2_**). Scale bars: 100 µm in (**c**) and 25 µm in (**d**).

## Discussion

3D-IHC reveals the spatial organization of molecular and cellular assemblies in the context of the tissue architecture. Here, we reported POD-nAb/FT-GO 3D-IHC as a multiplex nAb-based IHC of 3D tissues. POD-nAbs, camelid nAbs fused with POD, enhanced immunolabeling depth and enabled sensitive detections by combined with our original fluorescent TSA system, FT-GO. Immunolabeling depth was further enhanced by tissue permeabilization with Sca*l*eA2 solution. For multiplex labeling in 3D tissues with POD-nAb/FT-GO reaction, POD activity was quenched with NaN_3_ prior to subsequent rounds of reaction.

POD-nAb/FT-GO 3D-IHC is a high-speed and high-sensitive nAb-based 3D-IHC. We adapted the 3D-IHC technique to mouse brain slices of 1-mm thickness and succeeded in visualizing somata and processes of neuronal and glial cells in the thick tissue within three days (Fig. 1, 5, 6). Quenching of POD activity with NaN_3_ allowed for multiplex immunolabeling of 3D tissues by POD-nAb/FT-GO reaction (Fig. 8). Our 3D-IHC using POD-nAb/FT-GO reaction demonstrated high specificity for detection of exogenously expressed EGFP (98.7%) and tdTomato (98.8%) in mouse brain of 1-mm thickness (Fig. 4). POD-nAb/FT-GO 3D-IHC can be also used to detect endogenous proteins within 3D tissues. We generated a POD-nAb against ITGAM, a marker for myeloid lineage cells, and successfully labeled microglial cells in 1-mm-thick mouse brain slices (Fig. 6). Using the immunoreagent, we further visualized the local microglial activation surrounding Aß plaques of an AD mouse model in thick tissues (Fig. 7). Importantly, POD-nAb/FT-GO reaction of 1-mm-thick brain slices yielded 9.0-and 6.8-fold signal amplification compared with EGFP fluorescence and a synthetic fluorophore conjugated nAb against GFP, repsectively (Fig. 5).

Camelid nAbs fused with POD, POD-nAbs, enhanced penetration depth of probes and enabled sensitive detection of targets of 3D tissues by combined with FT-GO signal amplification. NAbs, recombinant antigen binding fragments from single-chain antibodies in camelids and selachians, are immunoreagents with high specificity, selectivity and reproducibility^20^. The small sizes of POD-nAbs (∼60 kDa), which comprises approximately 40% of molecular weight of conventional IgG Abs (∼150 kDa), might lead to deep penetration into large-scale tissues. Unlike conventional IgG and IgY Abs which deposited in the periphery of brain slices, POD-nAbs penetrated deep inside brain tissues and showed a high staining homogeneity (Fig. 2). NAbs conjugated synthetic fluorophore (∼20 kDa) are much smaller than POD-nAbs. These nAbs has been used for immunolabeling in large-scale tissues, visualizing subcellular details with a reasonable signal noise ratio^3,24^. However, the relatively low sensitivity of direct immunodetection^35^ can impede high-throughput imaging of large-scale tissues. POD fused with camelid nAbs enabled a TSA reaction, FT-GO, within large-scale tissues, providing a dramatic increase in sensitivity compared with direct IF with a synthetic fluorophore conjugated nAb (Fig. 5). The applicability of POD-nAbs (P-RAN bodies) have been documented using fluorochromized tyramide substrates in cultured cells and tissue sections^36^. The specificity, sensitivity and scalability of POD-nAbs would make them versatile tools for histochemical analysis.

Tissue permeabilization is critical for staining homogeneity of 3D-IHC to enhance immunolabeling depth. Of tested four tissue permeabilization protocols, we implemented Sca*l*eS tissue permeabilization, an incubation Sca*l*eA2 solution for 24 hr at 37 °C, to POD-nAb/FT-GO 3D-IHC. Consistent with a previous study demonstrating the reversibility of Sca*l*eA2 treatment that allows for retrospective IHC^37^, mouse brains slices permeabilized with Sca*l*eA2 solution retained sufficient antigenicity for a GFP and an RFP POD-nAb (Supplementary Figure 1, 2). While POD-nAbs against both GFP and RFP showed limited staining homogeneity in untreated control brain slices (Fig. 3b, d), they penetrated deeper inside brain tissues and reacted with their antigens at the center of brain slices following to Sca*l*eS tissue permeabilization (Fig. 3e, f). Moreover, EGFP and tdTomato fluorescence was preserved in brain slices permeabilized with Sca*l*eA2 solution (Supplementary Figure 1b_2_, 2b_2_). In contrast, immunoreactivity of the GFP and RFP POD-nAbs, and fluorescence of EGFP and tdTomato were markedly reduced after tissue permeabilization by PACT and iDISCO protocols (Supplementary Figure 1e, f, 2e, f), indicating these two tissue permeabilization protocols have deleterious effects on antigenicity and protein structures. However, these results do not indicate that POD-nAb/FT-GO reaction is compatible with only Sca*l*eS tissue permeabilization. Indeed, mouse brain slices permeabilized with CUBIC-HistoVIsion^12^ preserved immunoreactivity for the GFP and RFP POD-nAbs (Supplementary Figure 1, 2). Effective tissue permeabilization that preserves molecular and structural integrity and enhances immunolabeling depth would expand applicability and scalability of POD-nAb/FT-GO 3D-IHC.

FT-GO, a fluorescent TSA reaction, of 3D tissues allows for high sensitivity of our 3D-IHC protocol using POD-nAbs. TSA reaction including FT-GO utilizes POD to yield high-density labeling of targets *in situ*^25,26^. TSA has been widely used in detection procedures for histochemical analysis of tissue sections such as IHC, ISH, electron microscopy and neuroanatomical tract tracing^27–31^. However, this signal amplification technology has rarely applied to histochemistry of 3D tissues^44^. Relatively large size of POD-conjugated Abs that specifically recognize primary Abs or haptenized nucleotides and/or the need for additional steps for POD-catalyzed deposition of tyramide substrates on tissues might prevent implementation of TSA systems to 3D biological tissues. In the present study, we succeeded in adopting a TSA reaction, FT-GO, to detection procedure of millimeter-thick mouse brain tissues. POD-nAb/FT-GO reactions provided a remarkable increase sensitivity in 3D tissues: POD-nAb/FT-GO 3D-IHC for GFP yielded 9.0-and 6.8-fold signal enhancement in 1-mm-thick mouse brain slices, compared with EGFP fluorescence and direct IF with a GFP nAb conjugated with a synthetic fluorophore (Fig. 5). Multiplex labeling with POD-nAb/FT-GO reaction was achieved by quenching activity of POD with NaN_3_ (Fig. 7). NaN_3_ is effective at quenching Ab-conjugated POD and less deleterious for antigenicity, antigen–antibody binding, and tissue integrity compared with other methods such as incubation with hydrogen peroxidase and low pH buffer, and heat-mediated removal of Ab-POD complex^35,54^.

### Limitations of the study

In the present study, we developed POD-nAb/FT-GO 3D-IHC as an nAb-based multiplexed IHC of millimeter-thick tissues. A major limitation of the method is the scaling of the 3D-IHC. Information about long-range connectivity is missing and incomplete in the millimeter-thick brain tissues^34,55–57^. Modulation of antigen–antibody binding by manipulation of temperature, pH, and ionic strength, and/or facilitation of Ab diffusion by effective tissue permeabilization and adopting a higher incubation temperature can enhance penetration depth of Abs and staining homogeneity^9^. Moreover, transcardial perfusion of nAbs and conventional IgG Abs permits whole-organ and whole-body immunolabeling by greatly reducing the traveling distance for Abs to reach their targets^3,6^. These techniques might help in improving the applicability and scalability of POD-nAb/FT-GO 3D-IHC to whole-organ and whole-body levels.

Another major limitation of our method is that POD-nAbs available for 3D-IHC using FT-GO reaction are quite scarce. POD-nAbs improve immunolabeling depth to increase staining homogeneity and allow for sensitive detection of targets by combined with a multiplex TSA reaction in 3D tissues. Moreover, nAbs are highly specific and selective and immunoreagents with excellent solubility, superior stability, and high reproducibility. However, in contrast to the thousands of conventional IgG Abs that have been generated over the past decades, only a handful of nAbs are available and effective at histochemical analysis. The accelerating deposition rate of nAb structure data and sequence information into the public repositories^58–60^ might expand POD-nAb repertoire available to our 3D-IHC system.

## Methods

### Animals

Male and female C57BL/6J (Nihon SLC), *App^NL-G-F/NL-G-F^*(RBRC06344, RIKEN BioResource Research Center)^51^ and PV-FGL mice^53^ mice were used. C57BL/6J and PV-FGL mice were used at 8-16 weeks old, and *App^NL-G-F/NL-G-F^*mice were sacrificed at 4.5-and 6-month-old. *App^NL-G-F/NL-G-F^* and PV-FGL mice were maintained in C57BL/6J background. All mice were housed in specific pathogen-free conditions under a 12/12 hr light/dark cycle (light: 08:00–20:00) at 20-25 °C and variable humidity with *ad libitum* access to food and water. All animal experiments were approved by the Institutional Animal Care and Use Committees of Juntendo University (Approval No. 2021245 and 2021246). All animal procedures were conducted in compliance with ARRIVE (Animal Research: Reporting In Vivo Experiments) guidelines.

### Tissue preparation

Mice were anesthetized with an intraperitoneal injection of sodium pentobarbital (200 mg/kg; P0776, Tokyo Chemical Industry) and perfused transcardially with 20 mL of ice-cold phosphate buffered saline (PBS), followed by the same volume of ice-cold 4% paraformaldehyde (PFA) (1.04005.1000, Merck Millipore) in 0.1 M phosphate buffer (PB; pH 7.4). Brains were removed and postfixed in the same fixative overnight at 4°C.

### Tissue slicing

Brains were embedded in 4% agar (01028-85, Nacalai Tesque) in PBS and mounted on a vibrating tissue slicer (Linear PRO7N, Dosaka EM). The brains were cut into slices of 500-µm or 1-mm thickness.

### Tissue sectioning

For re-sectioning to test Ab penetration, immunolabeled brain slices were cryoprotectd in 30% sucrose in 0.1 M PB at 4°C, embedded in OCT compound (4583, Sakura Finetek) and frozen in isopentane cooled with liquid nitrogen. The slices were mounted on sliding microtomes (2000R, Leica Biosystems or REM-710, Yamato Koki) equipped with an electro-freezing component (MC-802A, Yamato Koki) and low temperature circulator (CCA-1112A, EYELA), and cut perpendicularly into 40-µm-thick sections^44,46^.

Frozen brain sections were prepared as described previously^61^. The fixed brain were cryoprotected in 30% sucrose in 0.1 M PB at 4°C and stored until use. Coronal brain sections were cut at 40-µm thickness on the sliding microtomes as described above.

### Cell line

293T cells were obtained from RIKEN BioResource Research Center (RCB2202). The cells were cultured in Dulbecco’s modified Eagle’s medium (11965-092, Thermo Fisher Scientific) supplemented with 10% fetal bovine serum (173012, Sigma-Aldrich), 2 mM L-glutamine (25030-081, Thermo Fisher Scientific), 1× MEM Non-Essential Amino Acid (11140-050, Thermo Fisher Scientific) and 1× Penicillin-Streptomycin (15070-063, Thermo Fisher Scientific) in a 5% CO2, 95% humidity incubator at 37 °C. When the cells were reached 60-70% confluence, they were passaged with TrypLE Express (12605-010, Thermo Fisher Scientific). The cell line is not listed as misidentified cell line in the ICLAC register.

### POD-nAb vectors construction

For the construction of pCAG-GFP POD-nAb1-WPRE and pCAG-RFP POD-nAb6-WPRE, a posttranscriptional regulatory element of woodchuck hepatitis virus (WPRE) sequence was amplified from pCAGGs-ChR2-Venus^62^ (#15753, Addgene) by polymerase chain reaction (PCR) and cloned to NotI site of pCAGEN^63^ (#11160, Addgene) to yield pCAG-WPRE. To obtain DNA fragments containing the coding sequence of P-RAN-GFP1 and P-RAN-GFP6, pCMV-P-RAN-GFP1^36^ (#106408, Addgene) and pCMV-P-RAN-RFP6^36^ (#106411, Addgene) were digested with AscI, and blunted with Klenow fragment (2140A, TAKARA) before further digestion with EcoRI. The fragments were ligated into pCAG-WPRE through EcoRI/EcoRV sites, yielding pCAG GFP POD-nAb1-WPRE and pCAG-RFP POD-nAb6-WPRE.

pCAG-ITGAM POD-nAb-WPRE construct was created by replacing the GFP-nAb1 sequence of pCAG-GFP POD-nAb1-WPRE with an ITGAM nAb sequence. The coding sequence of an nAb against ITGAM (VHHDC13)^47^ was synthesized by Eurofins Genomics KK.

### Antibodies

To produce POD-nAbs, 293T cells were transfected with pCAG-GFP POD-nAb1-WPRE, pCAG-RFP POD-nAb6-WPRE and pCAG-ITGAM POD-nAb-WPRE vectors using Lipofectamine 3000 Reagent (L3000001, Thermo Fiher Scientific) according to manufacturer’s instructions. Three days after transfection, the culture supernatant was collected, centrifuged at 3,000 g for 5 min at 4 °C, filtrated through 0.45 µm PVDF filters (SLHVR33RS, Merck Millipore). The supernatant was stored at 4 °C with 0.05% thimerosal (21624-32, Nacalai Tesque) and used for immunostainings.

Primary Abs used were; guinea pig (Gp) polyclonal anti-DsRed (red fluorescent protein) Ab (4.0 µg/mL, DsRed-GP-Af360, Frontier Institute, RRID: AB_2571648), rabbit (Rb) polyclonal anti-GFP Ab (10 µg/mL, A-11122, Thermo Fisher Scientific, RRID: AB_221569), rat (Rt) monoclonal anti-GFP Ab (1:1,000, 04404-26, Nacalai Tesque, RRID: AB_2313652), chicken (Ch) polyclonal anti-Green Fluorescent Protein Ab (2.0 µg/mL, GFP-1020, Aves, RRID: AB_10000240), Alexa Fluor 647-conjugated anti-GFP nAb (1:500, Cheomotek, gb2AF647, RRID: AB_2827575), Alexa Fluor 647-conjugated goat (Gt) anti-HRP Ab (1:100, 123-605-021, Jackson Immuno Research, RRID: AB_2338967), Gt polyclonal anti-Iba1 Ab (1:1,000, 011-27991, FUJIFILM Wako Pure Chemical Corporation, RRID:AB_2935833), mouse (Ms) monoclonal anti-RFP Ab (1:500, 409 011, Synaptic Systems, RRID: AB_2800533), and Gt polyclonal anti-tdTomato Ab (1:200, AB8181-200, SICGEN, RRID: AB_2722750).

Secondary Abs used were; Alexa Fluor 647-conjugated Gt anti-Ch IgY (10 µg/ml, A-21449, Thermo Fisher Scientific, RRID: AB_2535866), Alexa Fluor 647-conjugated Gt anti-Gp IgG (10 µg/ml, A-21450, Thermo Fisher Scientific, RRID: AB_2735091), Alexa Fluor 647-conjugated donkey (Dk) anti-Gt IgG (10 µg/ml, A-21447, Thermo Fisher Scientific, RRID: AB_2535864), POD-conjugated F(ab’)2 fragment Dk anti-Gt IgG (1:200, 705-036-147, Jackson Immuno Research, RRID:AB_2340392), Alexa Fluor 647-conjugated Dk anti-Ms IgG (10 µg/ml, A-31571, Thermo Fisher Scientific, RRID: AB_162542), Alexa Fluor 647-conjugated Gt anti-Rb IgG (10 µg/ml, A-21245, Thermo Fisher Scientific, RRID: AB_2535813) and Alexa Fluor 647-conjugated Gt anti-Rt IgG (10 µg/ml, A-21247, Thermo Fisher Scientific, RRID: AB_141778).

### Tissue penetration tests

To assess the tissue penetration of POD-nAbs and conventional IgG or IgY Abs, 1-mm-thick mouse brain slices infected with AAV2/PHP.eB CAG-EGFP-WPRE or CAG-tdTomato-WPRE were reacted for 4 days with POD-nAb culture supernatants or conventional IgG or IgY Abs. 0.3% (v/v) Triton-X100 was added to the culture supernatants. Conventional IgG or IgY Abs were diluted in PBS containing 0.3% (v/v) Triton-X100, 0.12% λ-carrageenan (035-09693; Sigma-Aldrich) and 1% normal donkey serum (S30-100ML, Merck Millipore) (PBS-XCD). Following Ab reaction, the brain slices were washed twice for 1 hr in 0.3% (v/v) Triton-X100 in PBS (PBS-X), fixed in 4% PFA in 0.1 M PB for 2 hr. 40 µm-thick re-sections were prepared from the slices as above. Then, the re-sections were washed for 10 min twice in PBS-X and reacted overnight with Alexa Fluor 647-conjugated Gt anti-HRP Ab (for POD-nAbs) or Alexa Fluor 647-conjugated secondary Abs (for conventional IgG or IgY Abs) in PBS-XCD at 4 °C. After washing for 10 min twice in PBS-X, the re-sections were mounted onto glass slides (Superfrost micro slide glass APS-coated, APS-01, Matsunami Glass) and coverslipped with 50% glycerol, 2.5% 1,4-diazabicyclo[2.2.2]octane (049-25712, FUJIFILM Wako Pure Chemical Industries), and 0.02% NaN_3_ in PBS (pH 7.4). All incubations were performed at 20–25°C, except where noted.

To test Sca*l*eA2 treatment on POD-nAb penetration, 1-mm-thick brain slices with infections of AAV2/PHP.eB CAG-EGFP-WPRE or CAG-tdTomato-WPRE were incubated in Sca*l*eA2 solution (4M urea, 0.1% [w/v] Triton X-100, 10% [w/v] glycerol) (21878933_Hama et al., 2011) for 24 hr at 37 °C. Sca*l*eA2-treated and PBS-stored brain slices were incubated in POD-nAb supernatants containing 0.3% (v/v) Triton-X100 or POD-nAb supernatants diluted with PBS-XCD for 24 hr, and washed twice for 1 hr in PBS-X at 20–25°C. After fixation in 4% PFA in 0.1 M PB for 2 hr at 20–25°C, re-sections of 40-µm thickness were prepared from the brain slices, labeled with Alexa Fluor 647-conjugated Gt anti-HRP Ab and mounted on slides as above.

### POD-nAb immunoreactivities on permeabilized tissues

To assess POD-nAb immunoreactivities on brain tissues permeabilized for 3D-IHCs, 1-mm-thick mouse brain slices infected with AAV2/PHP.eB CAG-EGFP-WPRE or CAG-tdTomato-WPRE were treated with tissue permeabilization of AbSca*l*e, CUBIC-HistoVision, iDisco and PACT. Tissue permeabilization procedures of each method followed the protocol as below. Considering 1-mm-thick slices, incubation time of each method was adjusted accordingly. Following tissue permeabilization, the brain slices were washed twice for 15 min in PBS, cryoprotected, embedded in OCT compound, frozen in cooled isopentane and cryosectioned on the freezing microtomes as above. Then, the re-sections were washed twice for 10 min in PBS-X and incubated overnight in POD-nAb supernatants containing 0.3% (v/v) Triton-X100. After washing twice for 10 min in PBS-X, the re-sections were reacted with Alexa Fluor 647-conjugated Gt anti-HRP and mounted onto slides. All incubations were performed at 20–25°C.

#### AbScale

Brain slices were incubated in Sca*l*eA2 solution for 24 hr at 37 °C^11^.

#### *CUBIC*-HistoVision

Brain slices were incubated in 50% CUBIC-L (T3740, Tokyo Chemical Industry) in distilled deionized water (DDW) for 2 hr and CUBIC-L for 24 hr at 37 °C^12^.

#### iDisco

After washing for 30 min in PBS, brain slices were serially incubated in 25%, 50%, 75% methanol (MeOH) in PBS, each for 30 min. The brain slices were incubated for 30 min in 100% MeOH and overnight in 66% dichloromethane (044-28305, FUJIFILM Wako Pure Chemical Industries) in MeOH. Then, the brain slices were washed twice for 15 min in MeOH, chilled on ice, and incubated overnight in 5% hydrogen peroxide in MeOH at 4°C. Following a wash in MeOH for 30 min, the brain slices were serially incubated in 75%, 50%, 25% MeOH in PBS, each for 30 min, washed twice for 15 min in PBS containing 0.2% (v/v) Triton-X100 (0.2% PBS-X) and incubated overnight at 37 °C in 0.2% PBS-X containing 20% Dimethyl sulfoxide (DMSO) and 0.3 M glycine. All incubations were performed at 20–25°C, except where noted^1,64^.

#### PACT

Brain slices were incubated in A4P0 hydrogel (4% acrylamide [161-0140, Bio-Rad] and 0.25% 2,2’-Azobis[2-(2-imidazolin-2-yl)propane]dihydrochloride [VA-044, FUJIFILM Wako Pure Chemical Industries] in PBS(–) [27575-31, Nacalai Tesque]) overnight at 4°C. The slices were then vacuum degassed for 10 min, placed under nitrogen for 10 min and incubated for 3 hr at 37 °C to initiate tissue-hydrogel hybridization. After washing twice in PBS(–) for 15 min at 20–25°C, the slices were incubated in 8% sodium dodecyl sulfate (31607-65, Nacalai Tesque) in PBS(–) for 24 hr at 37 °C. The slice were washed extensively overnight in PBS^39^.

### POD-nAb/FT-GO 3D-IHC

Brain slices of 500-µm or 1-mm thickness were incubated in Sca*l*eA2 solution or 24 hr at 37 °C, washed twice in PBS for 15 min and incubated for 20-24 hr in POD-nAb supernatants containing 0.3% (v/v) Triton-X100 at 20–25°C. The tissue permeabilization with Sca*l*eA2 was omitted from 3D immunolabeling with ITGAM POD-nAb. After washing four times in PBS-X for 30 min and thrice in 0.1 M PB for 5 min, the brain slices were incubated for 4 hr in an FT-GO reaction mixture that contained 10 µM CF488A (92171, Biotium), CF568 (92173, Biotium), CF640R (92175, Biotium) or CF647 tyramide (96022, Biotium), and 3 µg/mL glucose oxidase (16831-14, Nacalai Tesque) in 2% BSA in 0.1 M PB^35^. FT-GO reaction was initiated by adding 2 mg/mL of ß-D-glucose (049-31165, FUJIFILM Wako Pure Chemical Industries) into the reaction mixture and proceeded for 1-2 hr. The brain slices were washed twice in PBS-X for 15 min. Brain slices of *App^NL-G-F/NL-G-F^* mice were further incubated in 20 µg/mL 1-Fluoro-2,5-bis(3-carboxy-4-hydroxystyryl)benzene^52^ (FSB, F308, Dojindo) in PBS-X for 20-24 hr at 20– 25°C to label Aß plaques in the slices.

For 3D image acquisition, the brain slices were incubated in Sca*l*eS4 solution [4M urea, 40% (w/v) D-(–)-sorbitol, 10% (w/v) glycerol, 0.2% (w/v) Triton X-100, 25% (v/v) DMSO]^11^ for 12 hr at 37°C and mounted in an imaging chamber^44–46^. For tissue section preparation, the brain slices were fixed in 4% PFA in 0.1M PB overnight at 4°C, cryoprotected in 30% sucrose in 0.1 M PB and cut perpendicularly into 40-µm-thick sections as above.

### ScaleSF tissue clearing

For observation of EGFP fluorescence in 1-mm-thick brain slices, brain slices infected with AAV2/PHP.eB CAG-EGFP-WPRE were cleared by Sca*l*eSF method^44–46^. Brain slices were permeabilized in Sca*l*eS0 solution [20% (w/v) D-(–)-sorbitol, 5% (w/v) glycerol, 1 mM methyl-b-cyclodextrin, 1 mM g-cyclodextrin, and 3% (v/v) DMSO in PBS(–)]^11^ for 2 hr at 37°C, washed twice in PBS(–) for 15 min and cleared in Sca*l*eS4 solution for 10 to 12 hr at 37°C. Cleared brain slices were mounted in the imaging chamber as above.

### 3D-IHC with a synthetic fluorophore conjugated nAb

Following to permeabilization with Sca*l*eA2 solution for 24 hr at 37°C and twice washes with PBS for 15 min, brain slices infected with AAV2/PHP.eB CAG-EGFP-WPRE were reacted with an Alexa Fluor 647-conjugated anti-GFP nAb (1:500, Cheomotek, gb2AF647, RRID: AB_2827575) in PBS-XCD for 20-24 hr at 20–25°C. The brain slices were washed twice in PBS for 15 min, cleared with Sca*l*eS4 solution and mounted in an imaging chamber as above.

### FT-GO IF in tissue sections

FT-GO IF in free-floating sections was carried out as described previously^35^. In brief, tissues sections were quenched with 1% H_2_O_2_ in PBS and reacted with pAb diluted in PBS-XCD. After washing with PBS-X, the sections were reacted with POD-conjugated sAb, washed and incubated in an FT-GO reaction mixture. FT-GO reaction was initiated as described above and proceeded for 30 min. For IF in tissue sections stained by POD-nAb/FT-GO 3D-IHC, nAb-fused POD was quenched by an incubation in 2% NaN_3_ in PBS for 4 hr prior to the application of pAbs.

### AAV vector construction, production and injection

pAAV2-CAG-EGFP-WPRE and CAG-tdTomato-WPRE were constructed as follows. pCAGEN was digested with SalI, filled in with Klenow fragment, and digested with EcoRI to obtain a DNA fragment containing CAG promoter. The fragment was ligated into pAAV-MCS (Stratagene) which was digested with MluI and filled in with the Klenow fragment before further digestion with EcoRI to generate pAAV2-CAG-MCS. pCAG-WPRE was digested with HindIII, filled in with Klenow fragment, and digested with EcoRI to obtain a DNA fragment containing a WPRE sequence and a rabbit beta-globin polyadenylation (polyA) signal sequence. The DNA fragment was ligated into pAAV2-CAG-MCS which was digested with PmlI, filled in with Klenow fragment and digested with EcoRI to yield pAAV2-CAG-WPRE. The coding sequences of EGFP and tdTomato were amplified by PCR and ligated into pAAV2-CAG-WPRE through EcoRI/XhoI sites, yielding pAAV2-CAG-EGFP-WPRE and CAG-tdTomato-WPRE.

AAV vectors were produced and purified as follows^65^. pAAV2-CAG-EGFP-WPRE or pAAV2-CAG-tdTomato-WPRE and two helper plasmids, pHelper (Stratagene) and pBSIISK-Rep2-CapPHP.eB^66^, were transfected into 293T cells with polyethylenimine (23966, Polysciences). Virus particles were collected from the cell lysate and supernatant, purified by ultracentrifugation with OptiPrep (1114542, Axis-Shield) and concentrated by ultrafiltration with Amicon Ultra-15 (UFC905024, Merck Millipore). The physical titer of the virus vector (genome copies (gc)/mL) was determined with quantitative PCR using the purified virus solutions.

Intravenously administration of AAV vectors was performed as described previously^66^. After anesthetization with isoflurane (Pfizer) inhalation, AAV2/PHP.eB CAG-EGFP-WPRE (1.0 × 10^11^ gc) or CAG-tdTomato-WPRE (5.0 × 10^11^ gc) were injected into the retro-orbital sinus through a 29-or 30-gauge needle attached to a 0.5 mL syringe (SS-05M2913, TERUMO or 08-277, NIPRO).

### Image acquisition and processing

Images of tissue sections and cleared brain slices were acquired with a confocal laser scanning microscope (TCS SP8; Leica Microsystem). A 10× air (HCX PL APO 10x/0.40 CS, numerical aperture [NA] = 0.40, working distance [WD] = 2.20 mm, Leica Microsystems), 16× multi-immersion (HC FLUOTAR 16x/0.60 IMM CORR VISIR, NA = 0.60, WD = 2.5 mm, Leica Microsystems) and 25× water-immersion (HC FLUOTAR L 25x/0.95 W VISIR, NA = 0.95, WD = 2.50 mm, Leica Microsystems) objective lenses were used. Z-stack images were collected at 1.5 to 6.0 µm intervals at 512×512 or 1024×1024 pixel resolution. The confocal pinhole was adjusted to 1.0 or 1.5 Airy unit. Acquired images were tiled, stitched and processed to create MIP images using Leica Application Suite X software (LAS X, ver. 3.5.5.19976, Leica Microsystems) and ImageJ software^67^ (ver. 1.53, National Institutes of Health). The global brightness and contrast of the images were adjusted with ImageJ software.

## Quantification

To calculate the sensitivity and specificity of POD-nAb FT-GO/3D-IHC, FP or FT single-positive, and FP and FT double-positive neurons were counted in the cerebral cortex of re-sections using ImageJ software. Neurons were identified based on their morphological features:1) a round or ellipsoid-shaped cell body with smooth surface and 2) thick and tapering primary dendrites. The sensitivity was calculated by dividing the number of FP and FT double-positive neurons by that of FP positive neurons. The specificity was calculated by dividing the number of FP and FT double-positive neurons by that of FT positive neurons.

To assess the signal amplification resulting from POD-nAb/FT-GO reaction, the fluorescence intensity (arbitrary units [AU]) of cell bodies of labeled neurons was measured with ImageJ software. MIP images from a depth of 488 µm to 508 µm in the cerebral cortex were subjected to analysis. The identification of neurons was based on their morphological features.

## Statistics and Reproducibility

Three to five mice were used for each analysis. To establish the signal amplification effect of POD-nAb/FT-GO for EGFP fluorescence, the fluorescence intensity of POD-nAb/FT-GO reaction was compared to that of EGFP fluorescence in five mouse brains. The fluorescence intensity of four mouse brains stained by POD-nAb/FT-GO was compared to that stained with a synthetic fluorophore conjugated nAb against GFP to evaluate the signal amplification ability of POD-nAb/FT-GO for direct IF with an nAb. Statistical analyses were performed with the aid of GraphPad Prism 9 software (Version 9.4.1 (458), GraphPad Software). For data which had unequal variances, Welch’s t test was used for comparisons between groups. All tests were two-sided. Statistical significance was set at P < 0.05. Graphed data are represented as means ± standard deviations (SDs). The exact values of n are stated in the corresponding figure legends.

## Data availavility

The datasets generated during and/or analyzed during the current study and all biological materials reported in this article are available from the corresponding authors on reasonable request.

## Acknowledgements

We would like to thank Kisara Hoshino, Yoko Ishida, Ayaka Ansai, and Naoko Imai (Juntendo University) for their technical assistance. This study was supported by JSPS KAKENHI (JP20K07231 and JP23K06310 to K.Y.; JP20K07743 to M.K.; JP21H02592 and JP23K20044 to H.H.). This study was also supported by the Japan Agency for Medical Research and Development (AMED) (JP21dm0207112 to H.H.), Moonshot R&D from the Japan Science and Technology Agency (JST) (JPMJMS2024 to H.H.), Fusion Oriented Research for disruptive Science and Technology (FOREST) from JST (JPMJFR204D to H.H.).

## Author contributions

Conceptualization, K.Y.; Methodology, K.Y.; Validation, K.Y.; Investigation, K.Y.; Resources, K.Y.; Data Curation, K.Y.; Writing – Original Draft, K.Y,; Writing – Review & Editing, K.Y., M.K., and H.H.; Visualization, K.Y.; Supervision, H.H.; Project Administration, K.Y.; Funding Acquisition, K.Y., M.K., and H.H.

