## Supplementary Figure for "A Three Dimensional Immunolabeling Method with Peroxidase-fused Nanobodies and Fluorochromized Tyramide-Glucose Oxidase Signal Amplification"

Supplementary Figure 1. GFP1 POD-nAb immunoreactivity in brain slices processed for tissue permeabilization.

Supplementary Figure 2. RFP6 POD-nAb immunoreactivity in brain slices processed for tissue permeabilization.

Supplementary Figure 3. 3D-IHC with an Alexa Fluor 647-conjugated anti-GFP nAb..

Supplementary Figure 4. The detection of activated microglia with an ITGAM POD-nAb.

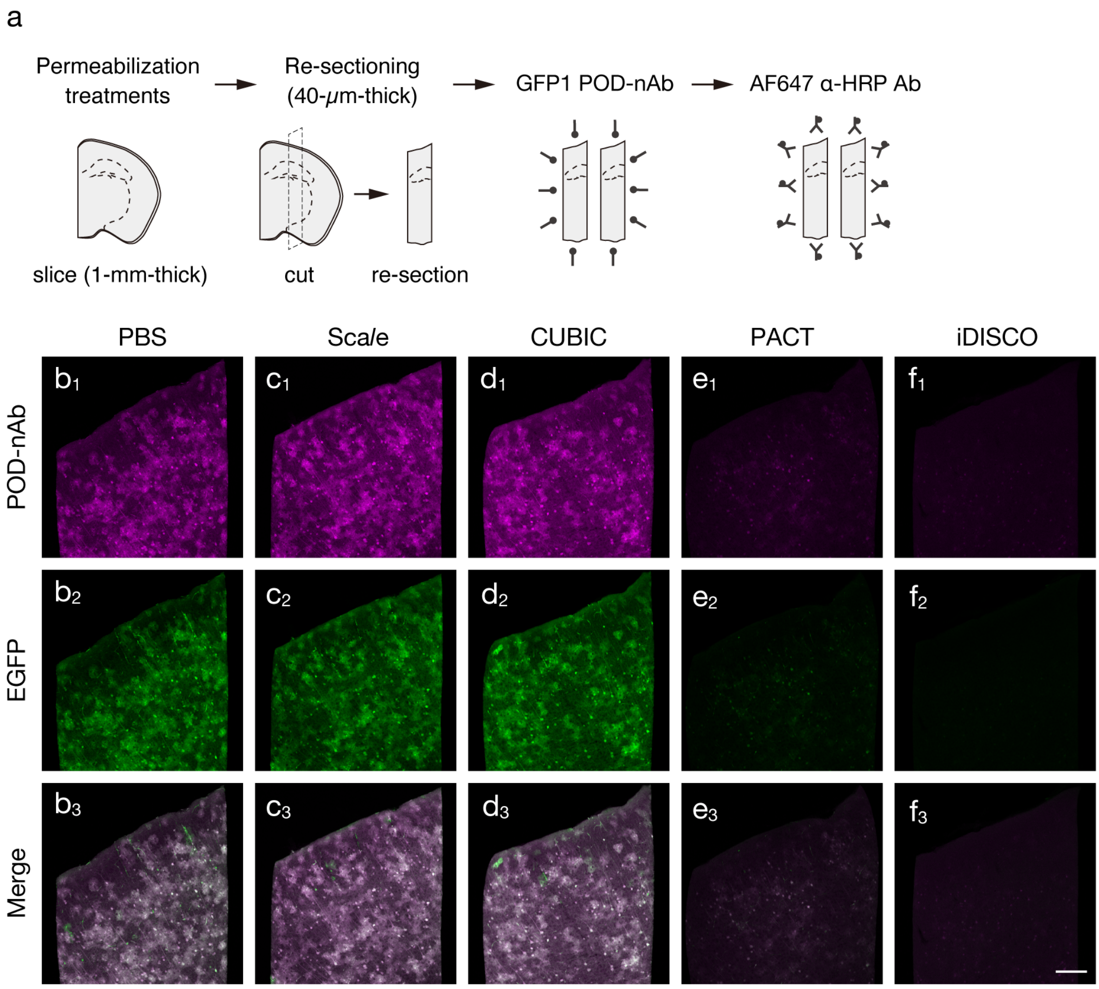


**Supplementary Figure 1| GFP1 POD-nAb immunoreactivity in brain slices processed for tissue permeabilization.**
**a**) Schematic diagram of an experimental procedure for testing tissue permeabilization on POD-nAb immunoreactivity. Brain slices are infected with AAV2/PHP.eB CAG-EGFP-WPRE. **b-f**) GFP1 POD-nAb IHC in mouse brain slices processed for control (**b**), Sca*l*e (**c**), CUBIC (**d**), PACT (**e**) and iDISCO (**f**) tissue permeabilization (n = 3 animals for each condition). $\mathbf{b}_{1,2}$, $\mathbf{c}_{1,2}$, $\mathbf{d}_{1,2}$, $\mathbf{e}_{1,2}$, $\mathbf{f}_{1,2}$) Representative images of immunoreactivity for GFP1 POD-nAb (magenta, $\mathbf{b}_1$, $\mathbf{c}_1$, $\mathbf{d}_1$, $\mathbf{e}_1$, $\mathbf{f}_1$) and EGFP fluorescence (green, $\mathbf{b}_2$, $\mathbf{c}_2$, $\mathbf{d}_2$, $\mathbf{e}_2$, $\mathbf{f}_2$). $\mathbf{b}_3$, $\mathbf{c}_3$, $\mathbf{d}_3$, $\mathbf{e}_3$, $\mathbf{f}_3$) Merged images of GFP1 POD-nAb immunoreactivity and EGFP fluorescence. Images are acquired with the same parameters for comparisons. AF: Alexa Fluor. Scale bar: 200 μm.

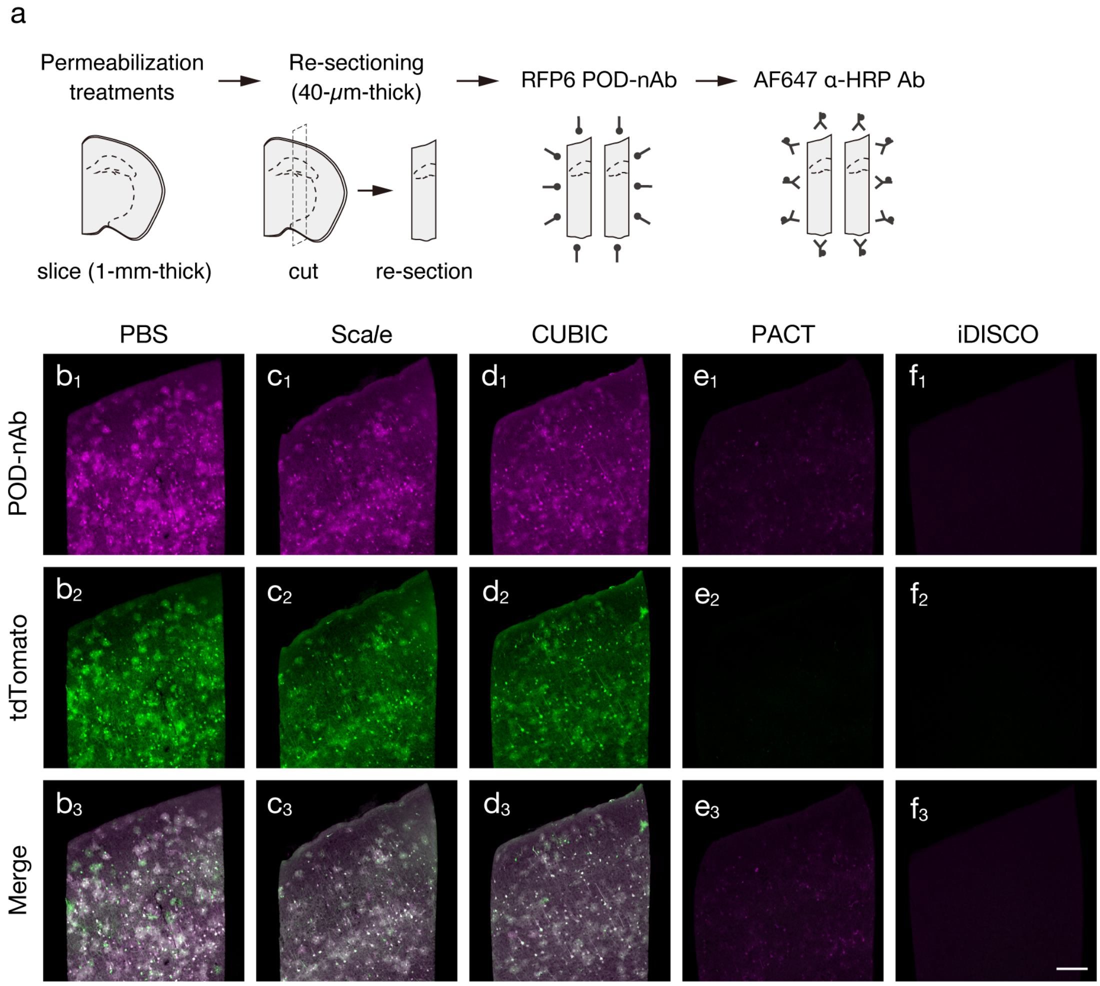


**Supplementary Figure 2| RFP6 POD-nAb immunoreactivity in brain slices processed for tissue permeabilization.**
**a**) Schematic diagram of an experimental procedure for testing tissue permeabilization on POD-nAb immunoreactivity. Brain slices are infected with AAV2/PHP.eB CAG-tdTomato-WPRE. **b-f**) RFP6 POD-nAb IHC in mouse brain slices processed for control (**b**), Sca*l*e (**c**), CUBIC (**d**), PACT (**e**) and iDISCO (**f**) tissue permeabilization (n = 3 animals for each condition). $\mathbf{b}_{1,2}$, $\mathbf{c}_{1,2}$, $\mathbf{d}_{1,2}$, $\mathbf{e}_{1,2}$, $\mathbf{f}_{1,2}$) Representative images of immunoreactivity for RFP6 POD-nAb (magenta, $\mathbf{b}_1$, $\mathbf{c}_1$, $\mathbf{d}_1$, $\mathbf{e}_1$, $\mathbf{f}_1$) and tdTomato fluorescence (green, $\mathbf{b}_2$, $\mathbf{c}_2$, $\mathbf{d}_2$, $\mathbf{e}_2$, $\mathbf{f}_2$). $\mathbf{b}_3$, $\mathbf{c}_3$, $\mathbf{d}_3$, $\mathbf{e}_3$, $\mathbf{f}_3$) Merged images of RFP6 POD-nAbs immunoreactivity and tdTomato fluorescence. Images are acquired with the same parameters for comparisons. AF: Alexa Fluor. Scale bar: 200 μm.

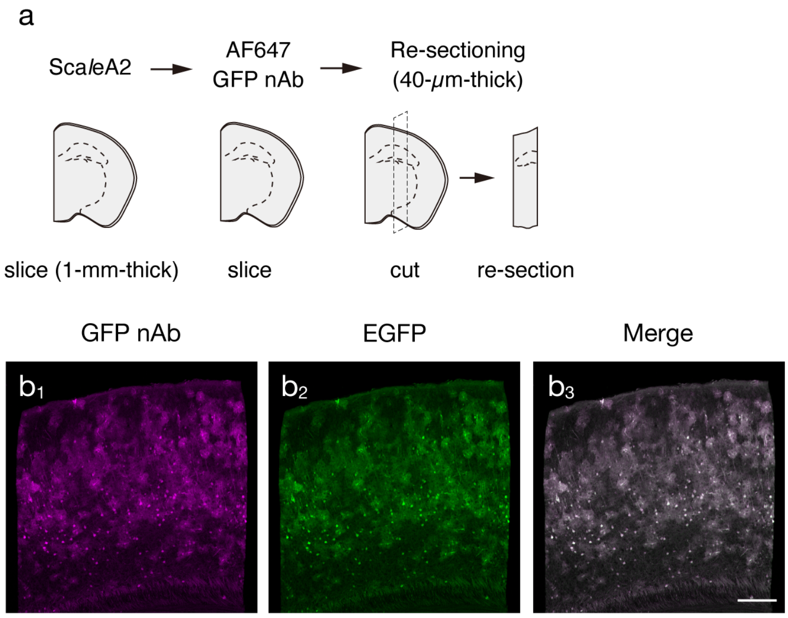


**Supplementary Figure 3| 3D-IHC with an Alexa Fluor 647-conjugated anti-GFP nAb.**

**a**) Schematic diagram of 3D-IHC with an Alexa Fluor 647-conjugated anti-GFP nAb and its analysis. Brain slices are infected with AAV2/PHP.eB CAG-EGFP-WPRE. **b**) GFP nAb immunoreactivity in a re-section prepared from a 1-mm-thick mouse brain slice processed for 3D-IHC (n = 3 animals). $\mathbf{b}_{1,2}$) Representative images of immunoreactivity for the GFP nAb (magenta, $\mathbf{b}_1$) and EGFP fluorescence (green, $\mathbf{b}_2$). $\mathbf{b}_3$) A merged image of ($\mathbf{b}_1$) and ($\mathbf{b}_2$). AF: Alexa Fluor. Scale bar: 200 μm.

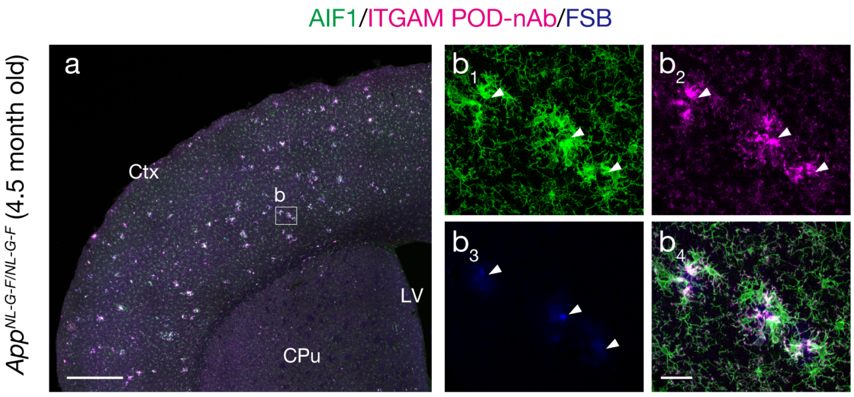


**Supplementary Figure 4| The detection of activated microglia with an ITGAM POD-nAb.**

**a**) An immunostaining with an AIF1 IgG Ab (green) and an ITGAM POD-nAb (magenta) in an $App^{NL\text{-}G\text{-}F/NL\text{-}G\text{-}F}$ brain section (n = 3 animals). The section is also fluorescently labeled with FSB (blue) for Aß plaque detection. **b**) A higher magnification image in a rectangle in (**a**). $\mathbf{b}_{1\text{-}3}$) Representative images of AIF1 IgG Ab immunoreactivity ($\mathbf{b}_1$), ITGAM POD-nAb immunoreactivity ($\mathbf{b}_2$) and FSB labeling ($\mathbf{b}_3$). $\mathbf{b}_4$) A merged image of ($\mathbf{b}_1$), ($\mathbf{b}_2$) and ($\mathbf{b}_3$). Arrowheads indicate the positions of Aß plaques labeled with FSB. CPu: caudate-putamen, Ctx: cerebral cortex, LV: lateral ventricle. Scale bar: 500 µm in (**a**) and 30 µm in (**b**).
